# Overexpression of human alpha-Synuclein leads to dysregulated microbiome/metabolites with ageing in a rat model of Parkinson disease

**DOI:** 10.1101/2020.12.23.424226

**Authors:** Yogesh Singh, Christoph Trautwein, Joan Romani, Madhuri S Salker, Peter H Neckel, Isabel Fraccaroli, Mahkameh Abeditashi, Nils Woerner, Jakob Admard, Achal Dhariwal, Morten S Dueholm, Karl-Herbert Schäfer, Florian Lang, Daniel Otzen, Hilal A Lashuel, Olaf Riess, Nicolas Casadei

## Abstract

Since Braak’s hypothesis stating that sporadic Parkinson’s disease follows a specific progression of the pathology from the peripheral to the central nervous system and can be monitored by detecting accumulation of the alpha-Synuclein protein. There is growing interest in understanding how the gut (commensal) microbiome can regulate alpha-Synuclein accumulation which can lead to PD. We studied a transgenic rat model overexpressing the human alpha-Synuclein and found that the protein overexpression resulted in gut alpha-Synuclein expression and aggregation in the gut neurons with advancing age. A progressive gut microbial composition alteration characterized by the reduction of Firmicutes to Bacteroidetes ratio could be detected in the young transgenic rat model and interestingly this ratio was then increased with aging. This observation was accompanied in older animals by intestinal inflammation, increase gut permeability and a robust alteration in metabolites production characterized by the increase of succinate level in the feces and serum. Manipulation of the gut bacteria by short-term antibiotics treatment revealed a complete loss of short-chain fatty acids (SCFAs) and reduction in succinate levels. Although antibiotics treatment did not change alpha-synuclein expression in the enteric nervous system of the colon, it can reduce alpha-synuclein expression in the olfactory bulb of the transgenic rats. In summary, synchronous with ageing, our data emphasize that the gut microbiome dysbiosis leads to a specific alteration of gut metabolites which are reflected in the serum and can be modulated by the environment.

## Introduction

The traditional hallmark of Parkinson’s disease (PD) is the presence of Lewy bodies (LBs), Lewy neurites (LNs) and the loss of dopaminergic (DA) neurons within the *substantia nigra pars compacta* (*SNpc*)^1–4^. LBs and LNs are structure resulting from the misfolding and aggregation of the alpha-Synuclein (α-Syn) protein^4^. Exact mechanisms leading to the aggregation of α-Syn and contributing to its toxicity are still unclear. Notably, the dopaminergic neurons of the *SNpc* appear particularly vulnerable to the effects of α-Syn aggregates and correlating to the clinical phenotype^4^. Surprisingly, new data has revealed that LBs and LNs are also found in the periphery including in nerves of the gastrointestinal tract (GIT)^5^.

Although PD is classically classified as a disease of the central nervous system, but numerous observations suggest that PD is not a single disease^6^ and that in specific form of PD is originating from peripheral tissues such as the gut epithelial cells called enteroendocrine cells (EECs), neurons of the GIT and olfactory system^2,7–9^. It is suggested that amyloidogenic α-Syn seeds may transfer from neuron to neuron as a result of retrograde transport causing accumulation of α-Syn in the bowel^2,10–13^. At the early stages of the disease, PD patients frequently exhibit non-motor symptoms such as olfactory malfunction, constipation and depression^14^. GIT dysfunction (in particular constipation) is observed in approximately 80% PD patients (sporadic and familial forms) before the occurrence of motor symptoms^14,15^. Recent patient study indicating two subtypes of PD, (1) brain-to-gut and (2) gut-to-brain with respect to α-Syn pathology, thus, clarifies the issue with different potentially conflicting earlier report in the PD field^16^.

Several studies have highlighted an association of gut microbiome dysbiosis in PD patients compared with healthy controls^1,17,18^. Whilst, individual bacterial composition varied from different PD-cohorts, only a few bacterial genera appeared to be consistent among different studies (e.g. *Bifidobacterium, Akkermansia and Lactobacillus*)^19,20^. Some were conflicting, in particular the *Lactobacillus* genus^21^ with some findings suggesting higher abundance^17,22–26^ whilst others suggested a lower abundance of *Lactobacillus*^18,27,28^. However, the major limitation of these studies is that the (PD patient’s) microbiome was quantified when the disease was already established. Further, it is not also known whether these changes in gut microbiome described play a causative role in the pathogenesis or are merely a consequence of disease^5^. Studying the microbiome before disease onset in humans is not possible due to unavailability of any known pre-clinical markers. Thus, rodent models are critical in understanding the physiological role of the gut microbiome on the disease development^29^.

In this context, the presence of synucleinopathy in the autonomic nervous system, accumulation of α-Syn in the bowel as well as altered intestinal microbiota correlated to motor symptoms pointing to a potential origin of synucleinopathy in the gut^30^. Gut bacteria can control the differentiation and function of immune cells in the intestine, periphery and in the brain^31–33^. A murine model that overexpressed the human α-Syn under the thy1 promoter (Thy1-α-Syn) suggested that gut microbiome transplants from PD patients into these mice induced parkinsonian-like motor dysfunction and that the microbiome was necessary to induce α-Syn pathology, neuroinflammation and motor defects^29^. Furthermore, using germ-free (GF) environmental conditions, this study showed that the gut microbiome was able to produce short-chain fatty acids (SCFAs) which were upregulated in the Thy1-α-Syn mice^29^. These SCFAs, especially butyrate, protect against intestinal hyperpermeability and decrease inflammation through modulation of microglial activation^34^. Additionally, SCFA-producing bacteria are less abundant in PD patients^35^. However, the exact role of SCFAs in neurodegeneration required further investigation due to conflicting results in murine and PD patient. Interestingly, antibiotic treatment in Thy1-α-Syn mice decreased the fine and gross motor function as well as gut motility defects highlighting that microbial signals modulate the α-Syn-dependent pathology^29^, thus providing a pivotal link between the gut microbiome and PD pathology. This finding suggests a potential influence of genetics on gut-microbiome interactions and synucleinopathies^29^. However, how a specific gut bacterial genera/species interacts with its host genetics with progressive ageing leading to PD like disease progression remains unknown.

Therefore, we hypothesized that α-Syn overexpression could be involved in modulating the gut microbiome composition and metabolite production which results in the induction of α-Syn misfolding and aggregation in enteric neurons. Accordingly, studying the microbiome dynamics, GIT physiology and function could lead to vital new insights into PD pathology. Given the broad pathophysiological effects of PD, we have used a previously described rat model overexpressing full-length non-mutated human α-Syn which includes the entire *SNCA* sequence with upstream regulatory promoter sequences and a flanking downstream region in a BAC construct^36^. Further, the rat BAC-SNCA transgenic PD (referred to as TG) model was shown to present a strong overexpression of α-Syn in the brain and to also reproduce the formation of pathological form of α-Syn. These TG rats develop early changes in novelty-seeking, avoidance, locomotor decline and an early smell deficit before the progressive motor deficit^36^. The observed pathological changes were linked with severe loss of structural dopaminergic integrity^36,37^. Moreover, these TG rats also showed bidirectional, trans-synaptic parasympathetic and sympathetic propagation of α-Syn^38^. This TG rat model holds close resemblance to the human PD disease progression, therefore, this model appeared to be ideal for testing the hypothesis that α-Syn expression regulates the gut bacterial abundance and pathophysiology of synucleinopathies.

Herein, we studied longitudinally the gut microbiome dynamics with ageing and report that with ageing, overexpression of human α-Syn changes the gut microbiota composition, metabolites and inflammation. Together, these results suggest that the humanized-rat model supports the notion that the gut microbiome could shape the progression of PD pathogenesis.

## Results

### Microbiome dynamics in the α-Syn TG rats

To determine the effect of human α-Syn overexpression on the gut microbiome’s composition and diversity of the gut microbiome, we analysed faecal pellets from the colon of the wild-type (WT) and TG rats by 16S ribosomal RNA (16S rRNA) gene amplicon sequencing. We used homozygous TG and control WT littermate rats, which were obtained from heterozygous mothers (Fig. 1a; described in Materials and Methods section) to negate any maternal (genetic) effects on microbiome analysis. The total numbers of reads between both WT and TG rat samples for each respective age group was shown for our sequencing depth (Suppl. Fig. 1a). Alpha diversity captures both the organismal richness of a sample and the evenness of the organisms’ abundance distribution^39,40^. We estimated the population diversity of the microbial community (alpha-diversity) by Chao 1 and Shannon-Weaver index with MicrobiomeAnalyst tool^41^ and MEGAN-CE software^42^ at phylum level. The alpha diversity Shannon-Weaver index in TG rats tended to be lower at each time point at phylum level, a trend, however, not reaching significance (Suppl. Fig. 1b). Most of the bacteria belonged to the Firmicutes and Bacteroidetes phyla and, both are typically the dominant phyla in the gut microbiome in humans and animals (Fig. 1b). At each age group, only a few bacterial phyla were significantly altered (WT vs TG comparisons), but most significant changes were observed during 1M (Proteobacteria; p=0.04), 2M (Aquificae; p=0.007) 2.5M (Firmicutes; p=0.02, Bacteroidetes; p=0.005, Actinobacteria; p=0.04 and Cyanobacteria; p=0.03), 6M (Actinobacteria; p=0.02, Verrucomicrobia; p=0.01 and Chloroflexi;p=0.01) and >12M (12-14M) (Bacteroidetes; p=0.02) of age in TG rats (Fig. 1b and Suppl. Fig. 1c). Further, with ageing the Bacteroidetes phylum was significantly reduced in TG rats (Fig. 1b and Suppl. Fig. 1c). The Proteobacteria phylum abundance was increased in >12M TG rats, however, it did not reach a significant level (Fig. 1b). Further, the Firmicutes and Bacteroidetes (F/B) ratio was calculated for the respective age group in WT and TG rats. The 2.5M age group had a significant reduction in F/B ratio in TG group (Fig. 1c). In sharp contrast, this ratio was significantly increased at >12M old TG group (Fig. 1c). Further, we performed two-way analysis of variance analysis (Two-way ANOVA) to find a correlation between change in the F/B ratio affected by age and bacteria. Our analysis suggested that with age there is a change in F/B ratio but both factors independently are no correlated (Fig. 1c). Additionally, we analysed the alpha diversities (Shannon-Weaver index and Chao 1) (Suppl. Fig. 1d) and the bacterial composition at the genera level (Fig. 1d). Surprisingly, even at an early age (2.5M), we were able to detect bacterial dysbiosis in TG rats compared to WT control rats and observed that ageing accelerates bacterial dysbiosis dynamics significantly (Fig. 1d and Suppl. Fig. 1e and Suppl. Fig. 2). The most interesting correlation was between *Lactobacillus* and *Alistipes*. With ageing TG rats had decreased % abundance of *Lactobacillus* whereas % abundance of *Alistipes* genus was significantly increased (Fig. 1e, f). Two-way ANOVA suggested that in WT and TG rats, with age there is change in the dynamics of *Alistipes* and *Lactobacillus* (Fig. 1e & f). Thus, overall, bacterial composition at genera level was significantly different between WT and TG rat at any given age group (Suppl. Fig. 2).

**Fig.1.**
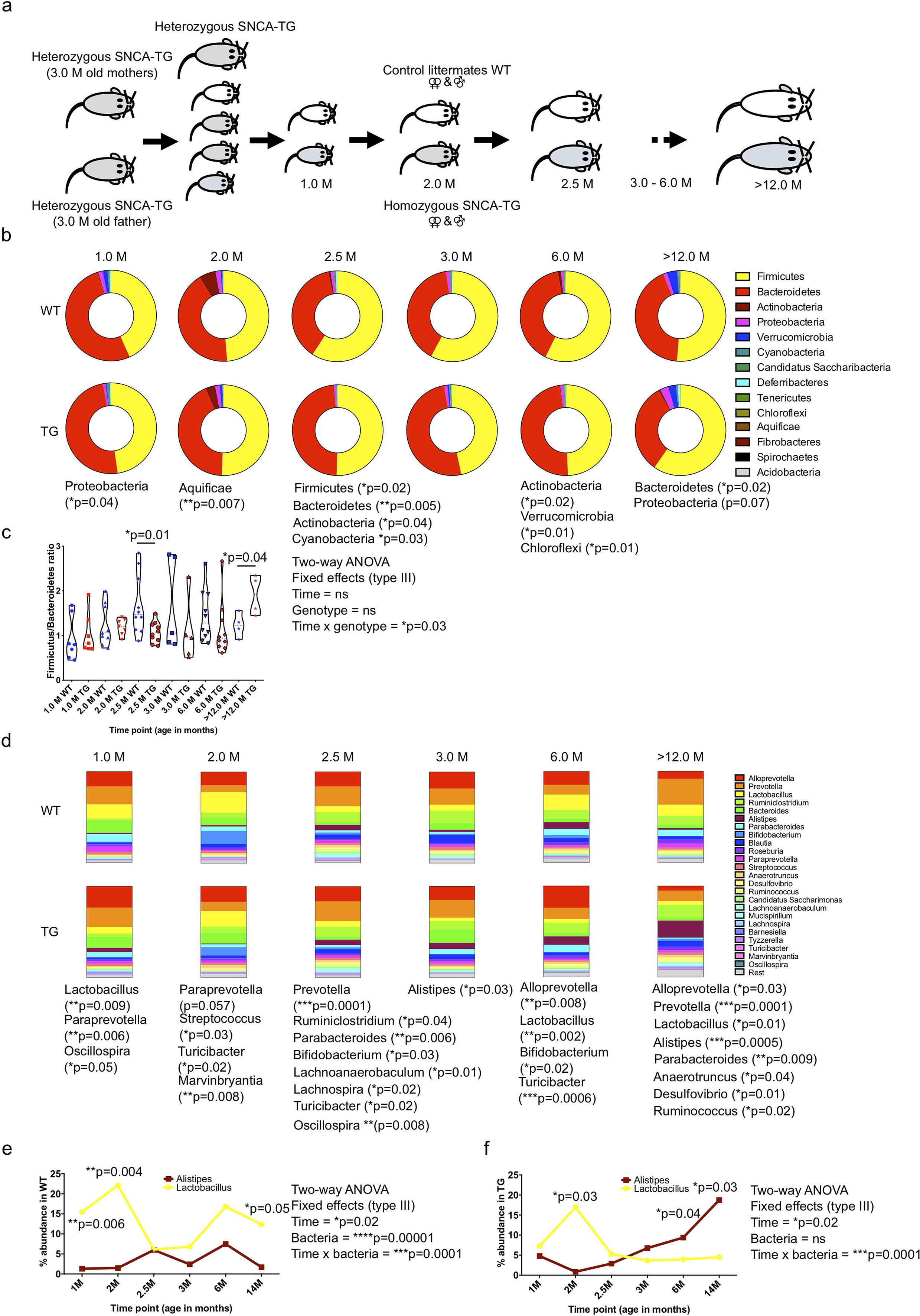
The gut microbiome dynamics with ageing, increased *Lactobacillus* and decreased *Alistipes* bacterial genera in the BAC-hSNCA TG rats. (a) Experimental design for the microbiome study. To understand the microbiome dynamics with ageing, heterozygous female and heterozygous male were used for breeding to keep the same gut microbiome from the mother to avoid maternal effect. When females were pregnant, male were removed from the cage and pubs were allowed to stay with the mothers for 3 weeks or until weaning. Three weeks old pubs were genotyped and then separated into either WT or homozygous BAC-SNCA transgenic groups (TG; refers to always homozygous until unless it is stated elsewhere) groups then faecal sample collections were started at the age of 1 month (4-5 weeks age) onwards (2, 2.5, 3, 6, >12 (12-14)) for the microbiome analysis by either 16s rRNA gene amplicon or shotgun sequencing methods. The animals were kept in 3-4 different cages to avoid the ‘cage affect’ bias in data analysis. Both male and females were used for the faecal sample collection depending on the breeding. (b) Representation of bacterial diversity (hollow pie chart) at phylum levels in WT and TG rats. (c) Firmicutes/Bacteroidetes (F/B) ratio was calculated for WT and TG rat samples and significant difference was observed at 2.5M and >12M age. F/B ratio was lower in TG rats at 2.5M age however, this ratio was reversed at >12M which was significantly higher in TG rats. Student’s t-test was performed for comparisons at a given age for WT and TG. Two-way ANOVA was performed to find the significance between different time points and genotypes. (d) The microbiome dynamics at genera level was estimated. Data are represented in colour-coded stacked-bar chart for each age group for the WT and TG rats. Each bar chart represents the % abundance of bacterial genera of the total bacteria in each group at particular age. Significant change in bacterial phyla is shown in asterisk for particular phyla together with p value signficance. (e,f) The dynamic representation of *Lactobacillus* and *Alistipes* with ageing in WT and TG respectively. A significant difference was observed with ageing in *Lactobacillus* and *Alistipes*. Two-way ANOVA was performed to find the significance between different time points and genotypes. P value significance represents *p≤0.05, **p≤0.01 and ***p≤0.001.

Several patients and murine model studies have suggested that *Lactobacillus* is abundantly present in relation to PD^17,23,24,43^. In contrast, other studies including ours suggested that *Lactobacillus* is decreased in PD models as well as in PD patients^18,28,44^. To understand the discrepancies in the *Lactobacillus* abundance with respect to α-Syn overexpression, we performed a mother to children microbiome follow up study. Microbiome composition analyses suggested that at 2M of age, WT and TG rats have a different pattern of *Lactobacillus sp*. in their feces (Suppl. Fig. 3a, b, & c). There was no detectable effect of the cages on the microbiome composition (Suppl. Fig. 3a).

Different environments as found in different animal facilities could change the gut microbiome. Therefore, both WT and TG rats were also kept in two different animal facilities (Facility I and Facility II) and tested whether bacterial composition was different or similar. We found that two different animal facilities did have a similar bacterial composition including *Lactobacillus* abundance (Suppl. Fig. 4a & b). However, *Lactobacillus sp*. abundance varied in TG rats, although the trend was less *Lactobacillus* compared with WT (Suppl. Fig 4b).

Next, we investigated whether keeping both genotype rats in the same cage, could change the gut microbiome of older TG rats (‘cage transfer transplant’) or whether ‘genetic factors’ were still predominant in determining the microbiome. Thus, we performed a pivotal experiment to answer this conjecture. Firstly, WT and TG (4 females only) rats were kept separately for >12M and at the end of the experimental period faecal pellets were collected (24 hours prior) (Suppl. Fig. 5a). Subsequently, the groups were then divided (2 WT and 2 TG) and kept together in fresh cage for one week and faecal pellets collected at the beginning (day 0) and end of the experiment (day 7). We found that *Lactobacillus* remained reduced whereas *Alistipes* continued to be elevated (Suppl. Fig. 5e & f). The ratio of *Lactobacillus* and *Alistipes* was not changed dramatically (before and after) due to the cage transfer experiment (Suppl. Fig. 5). Thus, in summary, α-Syn overexpression could have a key role in governing the expression of different bacterial genera including *Lactobacillus*.

### Metagenomic bacterial functional analysis in older TG rats

The microbial dysbiosis with ageing in TG rats (>12M) was observed by 16S rRNA gene amplicons sequencing. However, this method is limited as it relies on the amplification of a single gene using PCR-based of confined regions of the bacterial genome only. To increase the sensitivity of the microbiome sequencing, we performed shotgun sequencing which also allow to sequence the whole bacterial genome and also provides critical metabolic function interaction of the bacterial community. The shotgun sequencing depth and taxonomic rarefaction was shown in Suppl. Fig. 6 for both WT and TG rats. Again, the F/B ratio was significantly higher in TG compared with control WT littermates at phylum level (Fig. 2a, b). The alpha diversity was significantly reduced in TG rats (Fig. 2). Furthermore, based on genera taxonomic unit, we calculated the percentage (%) composition of the gut bacteria in WT and TG rats and found that *Lactobacillus* was reduced in >12M TG rats compared with WT, whereas *Alistipes* was significantly increased in the TG rats (Fig. 2c, d, e & Suppl. Fig. 6). Alpha and beta diversities represent the similarity (or difference) in organismal composition within and in between the samples. Alpha diversity was measured using the Shannon-Weaver index revealing that alpha-diversity was significantly higher in WT compared with TG rats (Fig. 2f). Next, using the principal component analysis (Bray-Curtis method) to investigate beta diversity, we found that the gut microbiome at the genus level from WT and TG rats cluster separately (Fig. 2g). Overall, our data indicate that α-Syn overexpression induces the gut microbiome dysbiosis in ageing TG rats.

**Fig. 2.**
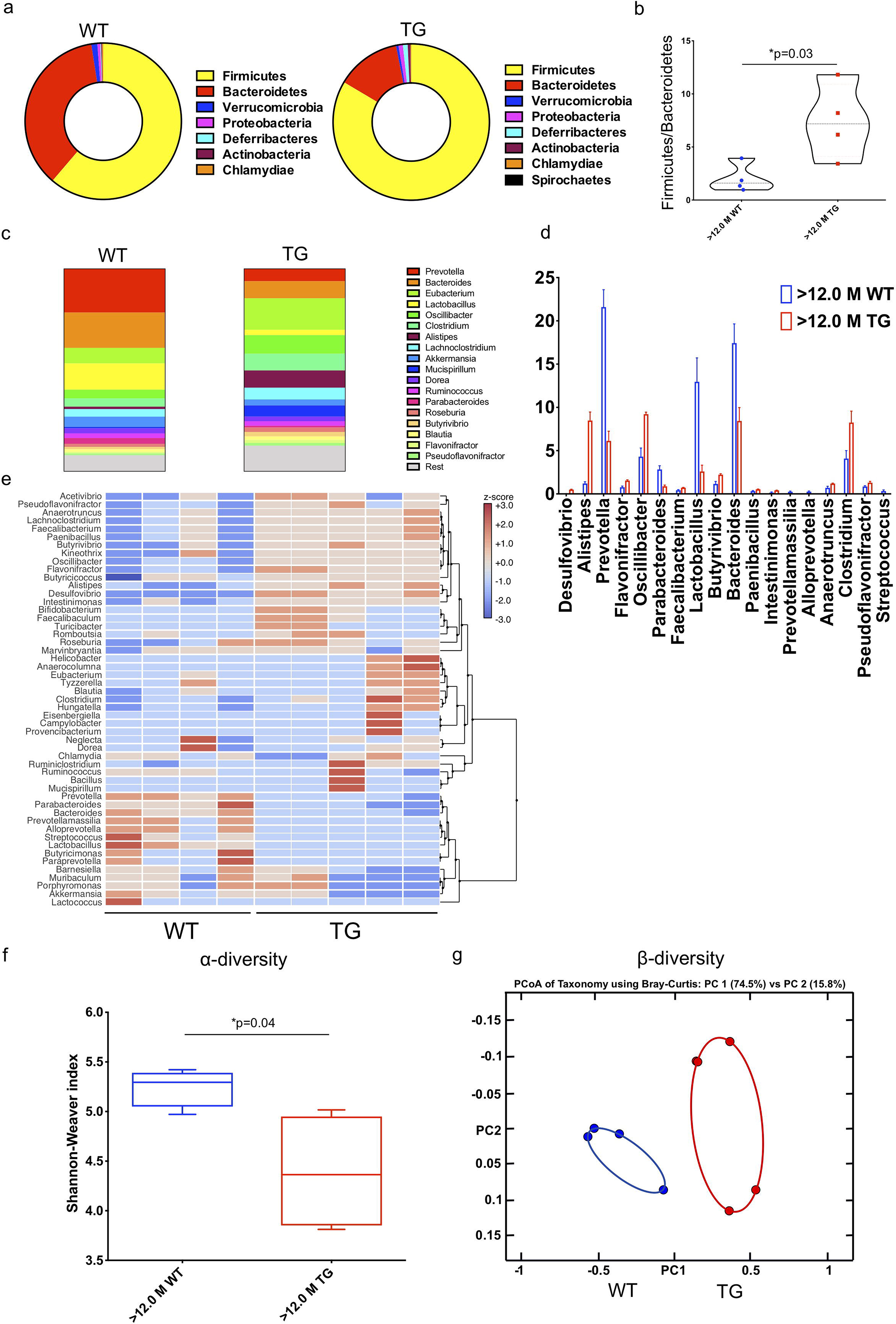
The gut dysbiosis in ageing TG rats. (a) To find out the bacterial dysbiotic bacterial strains, we have performed shotgun DNA sequencing and calculated the abundance of bacterial phyla (hollow pie charts) at >12M. (b) F/B ratio was significantly higher in TG compared with WT. (c, d) Several bacterial genera were significantly upregulated (*Desulfovibrio, Alistipes, Flavonifractor, Oscillibacter, Paenibacillus, Clostridium* and *Butyrivibrio*) and significantly downregulated (*Streptococcus, Bacteroidetes, Lactobacillus, Parabacteroidetes*, and *Prevotella*) at >12M age in TG rats. (e) Clustering of gut bacteria into three major clusters from WT and TG rats. (f) Alpha diversity at genera level was calculated using Shannon-Weaver index and found to be significantly lower diversity in TG rats compared with WT. (g) Beta diversity was estimated for both the genotypes of rats with Principal component analysis (PCoA) using Bray-Curtis method and both WT and TG rats cluster differently (PC1 54.5% and PC2 15.8). P value significance represents *p≤0.05.

### TG rats have a dysregulated metabolite production

To further understand change in the bacterial composition could also lead to different metabolites production, we performed proton nuclear magnetic resonance spectroscopy (^1^H-NMR)-based metabolomics from faecal and serum samples of younger (3M) and older (>12M) WT and TG rats (Fig. 3a). In total, we were able to quantify 31 different metabolites from feces and 38 metabolites from serum (Suppl. Fig. 7). Multivariate statistics using Partial Least Squares Discriminant Analysis (PLS-DA) was employed to identify differences between WT and TG rats for each age group (3M and >12M) for the feces (Suppl. Fig. 7a, b) and serum samples (Suppl. Fig. 7c, d). At >12M of age, there was a clear separation between WT and TG rat samples (Suppl. Fig. 7b, d), however this gap was less discrete for 3M age group samples (Suppl. Fig. 7a, c). When comparing variable importance in projection (VIP) scores of the PLS-DA feces analysis, we identified the most important metabolites - high levels of succinate in 3M TG rats whilst high levels of glutamate were discerned in >12M WT rats (Suppl. Fig. 7a, b). In serum, 3M WT rats showed elevated glucose levels, whilst >12M TG rats showed high lactate and succinate levels (Suppl. Fig. 7c, d).

**Fig. 3.**
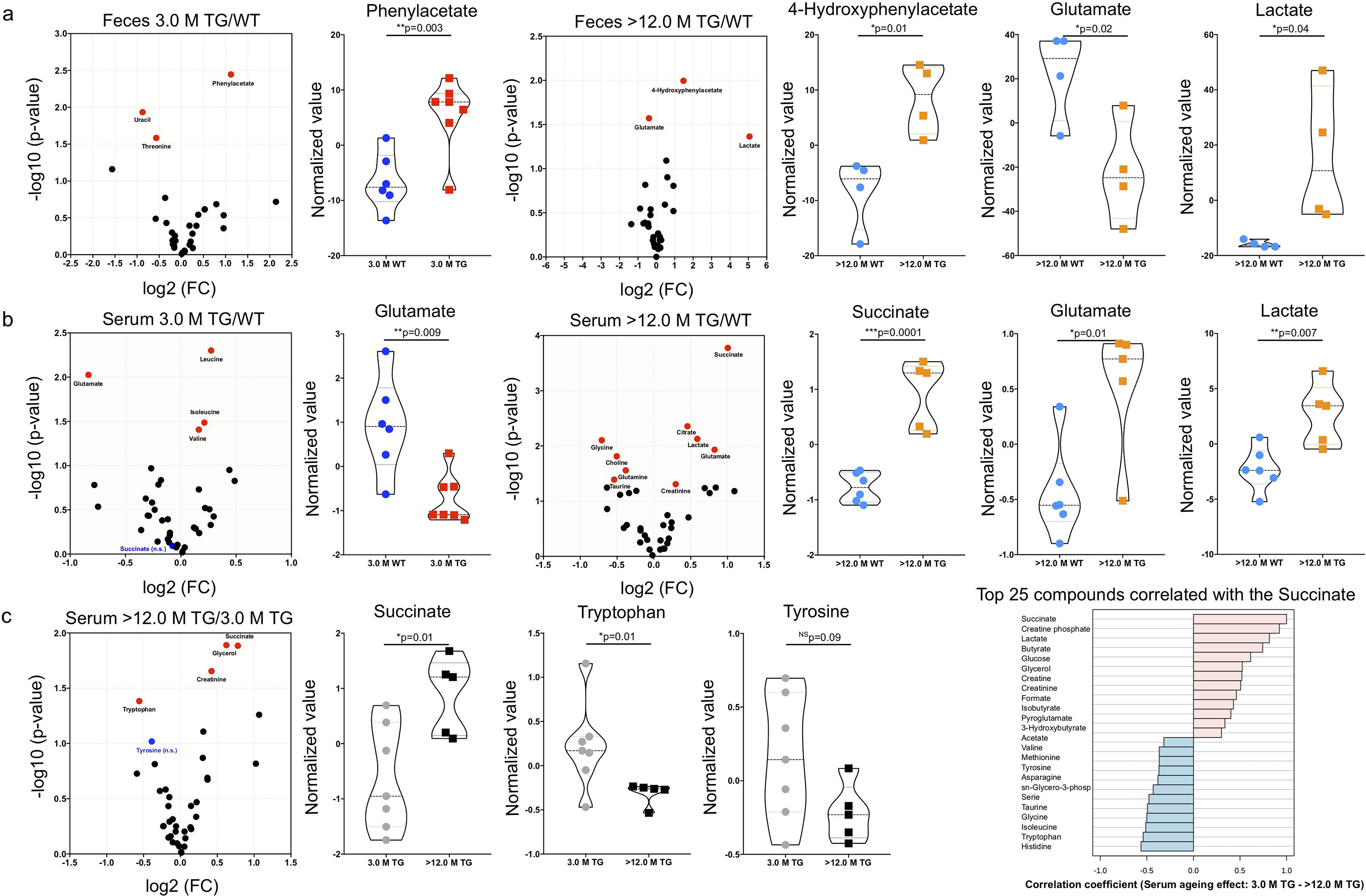
Ageing affects the succinate, tryptophan and tyrosine metabolites in the feces and serum. Metabolites combined fold change (FC > 1.2 (log2 FC)) and p-value < 0.05 (−log10 (p value)) volcano plot analysis identified effect of ageing between WT and TG rat feces and serum. (a) At 3M age feces genotype comparison TG/WT phenylacetate is significantly upregulated whereas at >12M age 4-Hydroxyphenylacetate and lactate were up while glutamate down. (b) In serum samples glutamate is significantly decreased at 3M whereas at >12M succinate, glutamate and lactate are increased. (c) When only observing the ageing effect on TG serum metabolites, succinate is significantly upregulated and succinate correlation coefficient analysis suggested that tryptophan and tyrosine were negatively correlated with succinate while other short chain fatty acids were positively correlated. Statistically significant levels showed in the respective volcano and violin plots. P value significance represents *p≤0.05, **p≤0.01 and ***p≤0.001.

To support the findings of multivariate statistics we applied a stringent univariate volcano plot analysis (combination of fold change and t-test). Hereby, we compared the metabolites levels based on genotype at each age group (3M and >12M) when comparisons were made for TG *versus* WT (TG/WT) (Fig. 3a, b) in the feces and serum respectively. Further, we then compared the levels of the metabolites for each genotype with age (3M to >12M) for WT (Suppl. Fig. 8a) and TG (Fig. 3c). We used FC > 1.2 (log2 FC) and p-value < 0.05 (−log10 (p value)) by default for all the metabolites and plotted against both values as volcano plots (Fig. 4 & Suppl. Fig. 8). At 3M of age, feces genotype comparison TG/WT suggested that phenylacetate was significantly upregulated whereas at >12M of age (TG/WT) a divergent set of metabolites such as 4-Hydroxyphenylacetate and lactate were upregulated while glutamate was significantly downregulated (Fig. 3a). We identified the most significant changes in the serum samples from >12M old TG and WT rats. We observed very high levels of lactate and succinate in TG compared to WT (Fig. 3b). Interestingly, when we explored serum metabolites, we found that glutamate was significantly downregulated at younger age (3M). In contrast, with ageing succinate, glutamate and lactate were significantly upregulated in older (>12M) TG rats compared to WT (TG/WT comparisons at a given age) (Fig. 3b). Further, we performed the ageing comparison for each genotype either WT or TG and found that serum succinate in TG rats was significantly upregulated (Fig. 3c; left). Succinate correlation coefficient analysis suggested that tryptophan and tyrosine were negatively correlated with succinate while other SCFAs were positively correlated (Fig. 3c; right). In WT serum succinate tended to decline with ageing, a trend, however, not reaching statistical significance (Suppl. Fig. 8a). Whilst observing the ageing effect for feces, the correlation analysis with succinate suggested tyrosine is negatively correlated, whilst 4-hydroxyphenylacetate was positively correlated in TG rats (Suppl. Fig. 8b). Overall, it appears that dysregulated levels of succinate, lactate and tyrosine could be linked with PD progression.

**Fig. 4.**
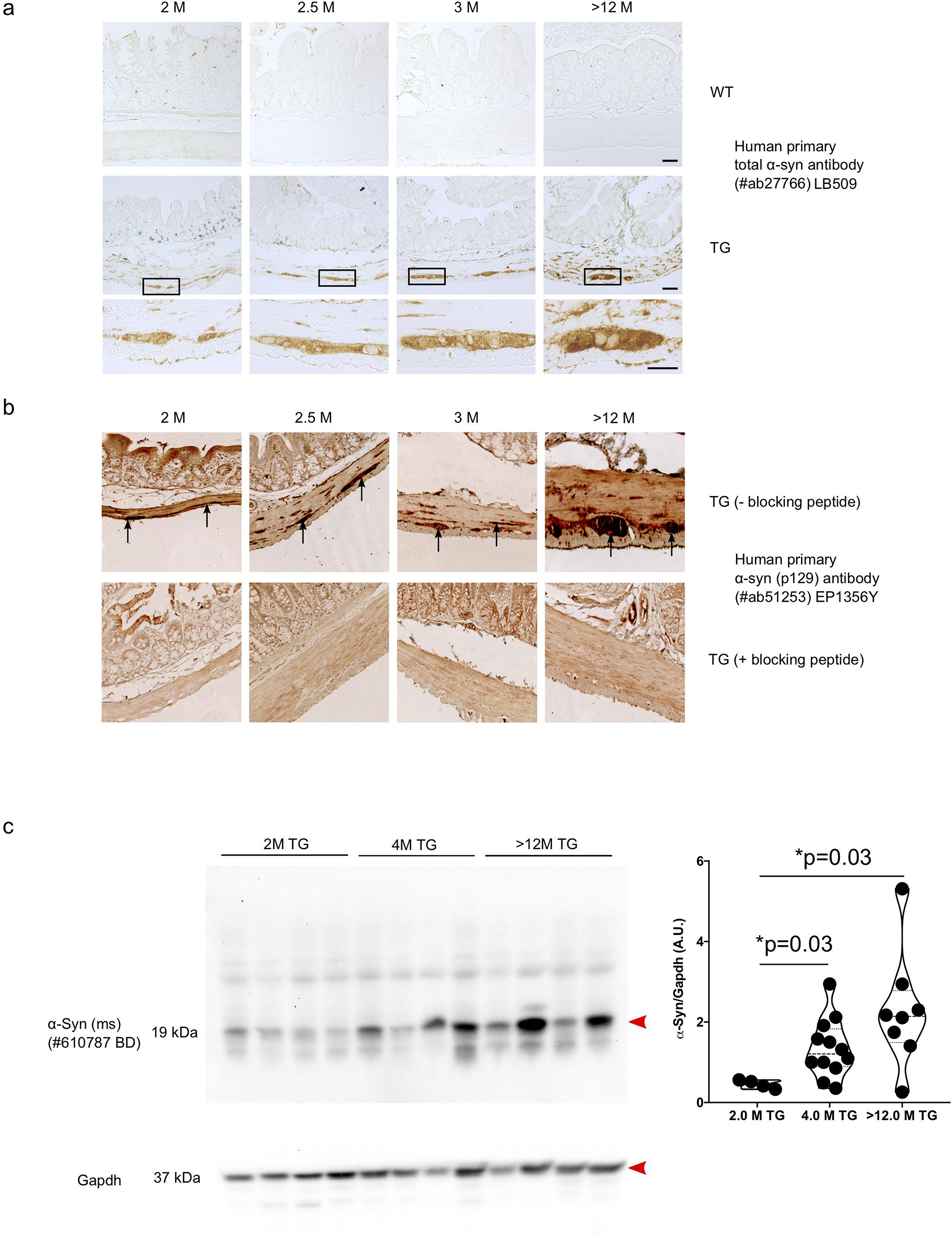
Accumulation of α-Syn in the colon of TG rats. (a) IHC was performed to identify the accumulation of total human α-Syn in 2, 2.5, 3 and >12M old WT and TG rats using human primary α-Syn antibody (LB509; #ab27766). With ageing accumulation of α-Syn was observed in TG rats whereas no positive staining was observed in WT rats as expected. (b) Further, pathological state of human α-Syn was examined in 2, 2.5, 3 and >12M in TG rats using human primary p-129 α-Syn antibody (EP1356Y; #ab51253). It appeared that pathological α-Syn was increased with ageing in TG rats. Further, p-129 α-Syn antibody was blocked using specific peptide for p-129 site for the antibody and no staining was observed suggesting the specificity of p-129 antibody. (c) One of the representatives immunoblot images of α-Syn and Gapdh staining’s are shown. We quantified the α-Syn amount using total α-Syn antibody (#BD 610787) and found that with ageing (2M) to 4M α-Syn expression in the colon tissues is significantly increased. Further ageing (4M to >12M) tended to affect the accumulation of α-Syn but not statistically significant difference was observed. P value significance represents *p≤0.05.

### Presence of synucleinopathy in the enteric nervous system (ENS) of TG rats

Using antibodies specific for human α-Syn, we detected using immunohistochemistry specific staining in the colon of TG rats and found presence of human α-Syn expression in the gut of TG rats whereas we did not detect any human α-Syn expression in WT rats (Fig. 4a). Human α-Syn staining was mostly found in the ENS of the colon and also in the myenteric plexus of TG rats. Presence of the transgenic human α-Syn protein was also validated by immunoblotting (Suppl. Fig. 9a). Expression of the human α-Syn expression was also detected in the myenteric plexus and in the colon of TG rats using the whole-mount immunostainings of the Tunica muscularis (Supp. Fig. 9b).

To investigate the presence of a pathological forms of α-Syn and an accumulation of α-Syn in TG rats, we first performed the histological analysis of the colon. We could detect reproducibly the presence of phosphorylation of human α-Syn at serine 129 position which increased with ageing (Fig. 4b). Presence of synucleinopathy was investigated using trypsin digestion to identify the insoluble forms of α-Syn. Colon sections of WT rats presented α-Syn staining with endogenous rat antibody, however, after trypsin treatment for 2 hours, this staining was lost in the myenteric plexus (Suppl. Fig. 10). Yet, in sharp contrast to WT rats, TG rats exhibited positive staining for total human α-Syn after trypsin digestion even in young rats suggesting that aggregation of α-Syn already occurs at two months of age (Suppl. Fig. 10). Total accumulation of α-Syn is increased with age in TG rats (2M) and reached a plateau at the age of 4M (Fig. 4c). At later stages (>12M), α-Syn still tended to increase slightly, but no significant change to 4M was observed (Fig. 4c). Together, our data suggest that human α-Syn accumulation occurs in the colon of young TG rats.

It was previously described that in the brain samples of TG rats there are several small sized fragments of α-Syn^36^. To test this, we performed immunoblotting of human α-Syn from colon of TG rats and identified several small-sized fragments of α-Syn in different age groups (5M, >12M) (Suppl. Fig. 11a). We found 3 major truncated fragments of human α-Syn protein which we characterized further using specific α-Syn antibodies targeted at N- or C-terminal epitopes (Suppl. Fig 11c). Truncated fragments did not arise from the boiling of the proteins as samples prepared in the presence of DSP (a cross-linker) led to the same results (Supp. Fig. 11b). Interestingly, α-Syn exhibited two N-terminal truncation fragments and one C-terminal truncation fragment (Suppl. Fig 11c). Thus, α-Syn expression in the colon revealed in addition to brain pathology that intestinal tissue is valuable in assessing PD progression in TG rats.

### Physiological functions of the TG rats dysregulated with ageing

The colon is a significant site of water and salt absorption and, during this process it desiccates the feces. The dysregulation in the gut microbiome and metabolites could influence the intestinal permeability. Therefore, we used an Ussing chamber to evaluate the function of intestinal permeability and Na^+^ uptake. To do so, we measured the transepithelial Na^+^ current (ENaC) in the colon epithelium. We measured the ENaC current in young (2M) and older (12M) rats of both the genotypes and found that 2M and 12M TG rats had a lower ENac current compared with WT rats respectively (Fig. 5a, b, c). This could suggest that TG rats may exhibit a dysregulation of the Na^+^ absorption in the colon. Further, we found that the transepithelial resistance tended to decrease with ageing, however no significant change was observed between WT and TG at any age group (Fig. 5c). Interestingly, the feces’ water content showed no difference between 12M WT and TG rats (data not shown). Hypoxanthine is negatively correlated with intestinal inflammation and permeability in inflammatory bowel disease^45^. We found that hypoxanthine is significantly increased in 14M WT compared with 3M WT rats but not in TG rats (between 3M and >12M). Hypoxanthine was not significant between TG and WT rats (Fig. 5d). Nonetheless, gut dysbiotic metabolites could affect the Na^+^ uptake or intestinal permeability in TG rats.

**Fig. 5.**
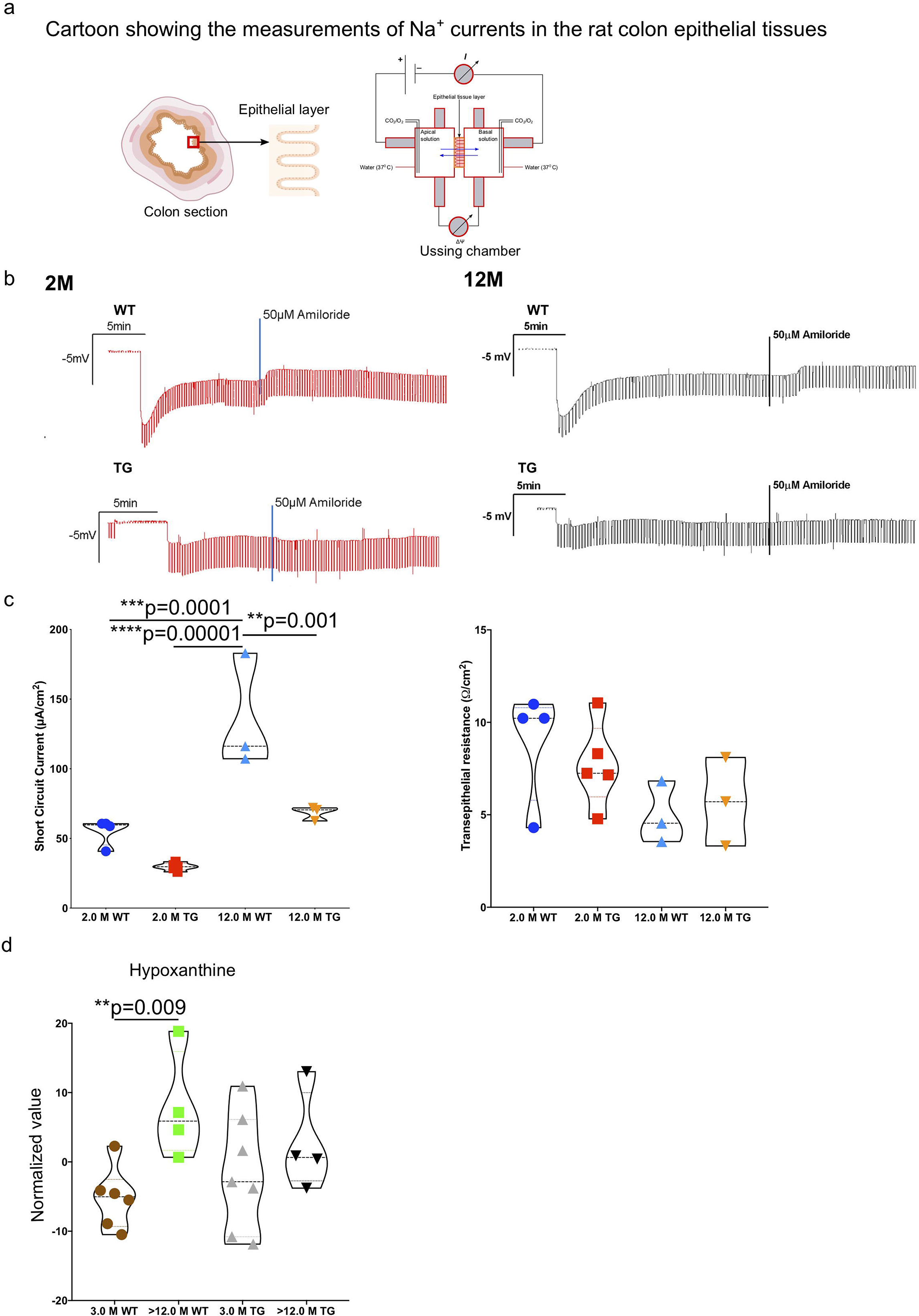
Increased permeability and transepithelial current in TG rats. (a) A schematic diagram showing the ‘Ussing chamber’ technique to measure the function of intestinal permeability and Na+ uptake to gut epithelial cells (Enac channel). (b) Original tracing illustrating the effect of currents (1 μA) and of amiloride (50 μM) on transepithelial potential across colonic epithelium from WT (upper) and TG (lower) from 2 and 12M rats. (c) Arithmetic mean +/− SEM (n=3-5/group) of amiloride-sensitive current across colonic epithelium (Na^+^ absorption) from TG and WT rats and resistance of colonic epithelium (permeability). (d) Faecal hypoxanthine levels measured by ^1^H-NMR from 3M and >12M WT and TG rats. P value significance represents *p≤0.05, **p≤0.01, ***p≤0.001 and ***p≤0.001.

### Increased local and systemic inflammation in TG rats

PD patients have a higher abundance of inflammatory proteins in the feces, possibly be due to dysbiotic changes in the gut microbiome^46–48^. Thus, we measured the faecal inflammatory proteins in TG rats. Faecal calprotectin level was significantly increased in older (8M) TG rats compared with WT, however no change occurred at an earlier age (3M) (Fig. 6a). To understand where the sources of these increased inflammatory proteins in the feces are, we extended our evaluation on the gene expression in the host intestine. Thus, we performed RNA-sequencing (RNA-seq) from the mucosa and submucosal tissues (defined as MS layer) from young (3M) and older (>12M) TG and control WT rats. Our RNA-seq experiments suggested that an enrichment of dysregulated genes involved in inflammation was detected with ageing in TG rats (Fig. 6b-e and Suppl. Fig. 12,13). Further, Ingenuity pathway analysis (IPA) suggested that several inflammatory pathways including innate and adaptive immunity were upregulated in >12M TG rats compared with 3M TG rats (Fig. 6b). Most interestingly, PD-1, PD-L1 cancer immunotherapy pathways, antioxidant action of vitamin C, RhoGDI signalling, Apelin cardiac fibroblast signalling pathway and LXR/RXR activation pathways were downregulated significantly in older TG (>12M) compared with 3M TG rats (Fig. 6b, c). More specifically, with ageing NFAT regulation of the immune response, cardiac hypertrophy signalling, neuroinflammation signalling pathway, G-beta gamma signalling, ephrin receptor signalling and opioid signalling pathway were upregulated with ageing in >12M TG rats compared to 3M TG rats (Fig. 6b-e). Additionally, whole genome wide analysis suggested that programmed cell death pathway genes were decreased, whilst immune system pathway genes were increased in old TG rats compared to young TG rats (Fig. 6b & Suppl. Fig. 13). Therefore, we tested the adaptive immune cells of younger (3M) and older (>12M) TG rats with their respective age matched WT controls. We found that ageing led to a significant increase in inflammatory IFN-γ cytokines from CD4^+^ T cells and also increased CD4^+^ T cells (Fig. 6f), whereas no significant difference was found at 3M age in between WT and TG (data not shown). Overall, data are suggesting that ageing TG rats have an activated inflammatory environment compared with younger TG rats.

**Fig. 6.**
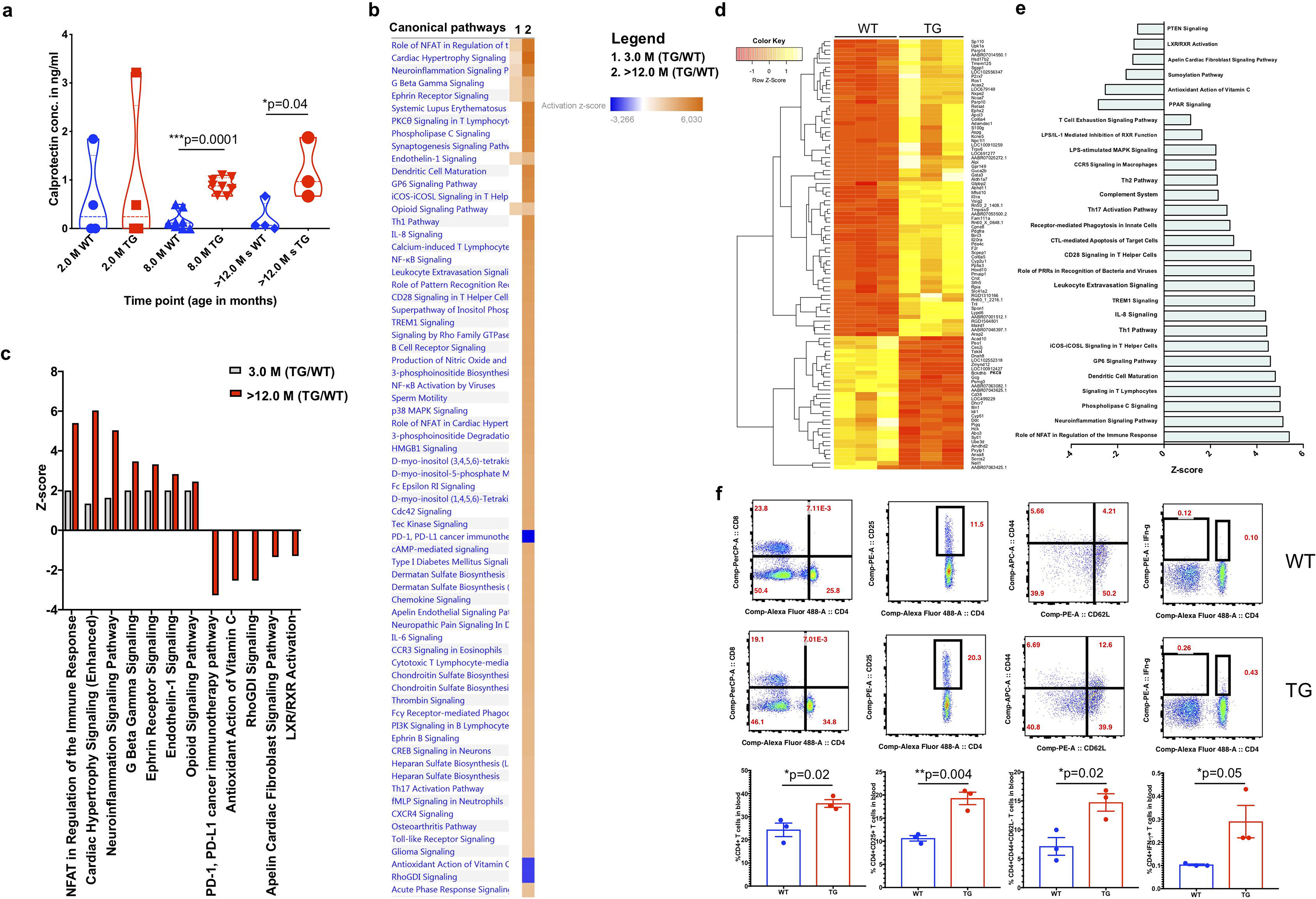
Increased inflammatory signals in the colon mucosa and submucosa (MS layer) of >12 M TG rats. (a) Fecal/serum calprotectin levels measured by ELISA in 2, 8 and >12 M WT and TG rats. (b) WT and TG (3M and >12M old) rat colon mucosa and submucosa (gut epithelium) was taken and subjected to RNA-seq. (b,c) Ingenuity pathway analysis (IPA) from TG and WT rats showed upregulation of several biological pathways in >12M TG rats. (d) The heatmap shows upregulation (yellow) and downregulation (red) genes in the gut MS layer. TG rats have higher expression of various genes involved in inflammation compared with WT. (e) IPA analysis in >12M TG rats suggested that several inflammatory pathways were changed. (f) FACS plots show increase in numbers of CD4^+^ T cells, CD4^+^CD25^+^ T cells (activated), CD^+^CD44^+^ T cells (memory) and CD4^+^ IFN-γ^+^ T cells were significantly higher in the blood of >12M TG rats compared with WT. Bar plots show the means +/− SEM. P value significance represents *p≤0.05.

### Antibiotics treatment affects the gut bacterial diversity, metabolites production and gene expression in the gut

To understand whether reducing the gut bacteria load could lead to changes in the inflammatory environment or α-Syn expression, we treated the WT and TG rats at 2.5M of age with broad-spectrum antibiotics for two weeks. Antibiotic treatment led to a reduction of live bacterial load and further reduced the amount of the total DNA obtained from an equal weight of the stool samples (Fig. 7a, b). The 16S rRNA amplicon sequencing suggested a tendency to reduce the F/B ratio and bacterial alpha diversity (phylum level) after antibiotics treatment for WT and TG rats respectively. However, no significant difference was observed (Fig. 7c-e). Beta diversity (PCoA of taxonomy using Bray-Curtis at phylum level) data suggested that antibiotics treated rats clustered into different groups compared with untreated rats (Fig. 7f).

**Fig. 7.**
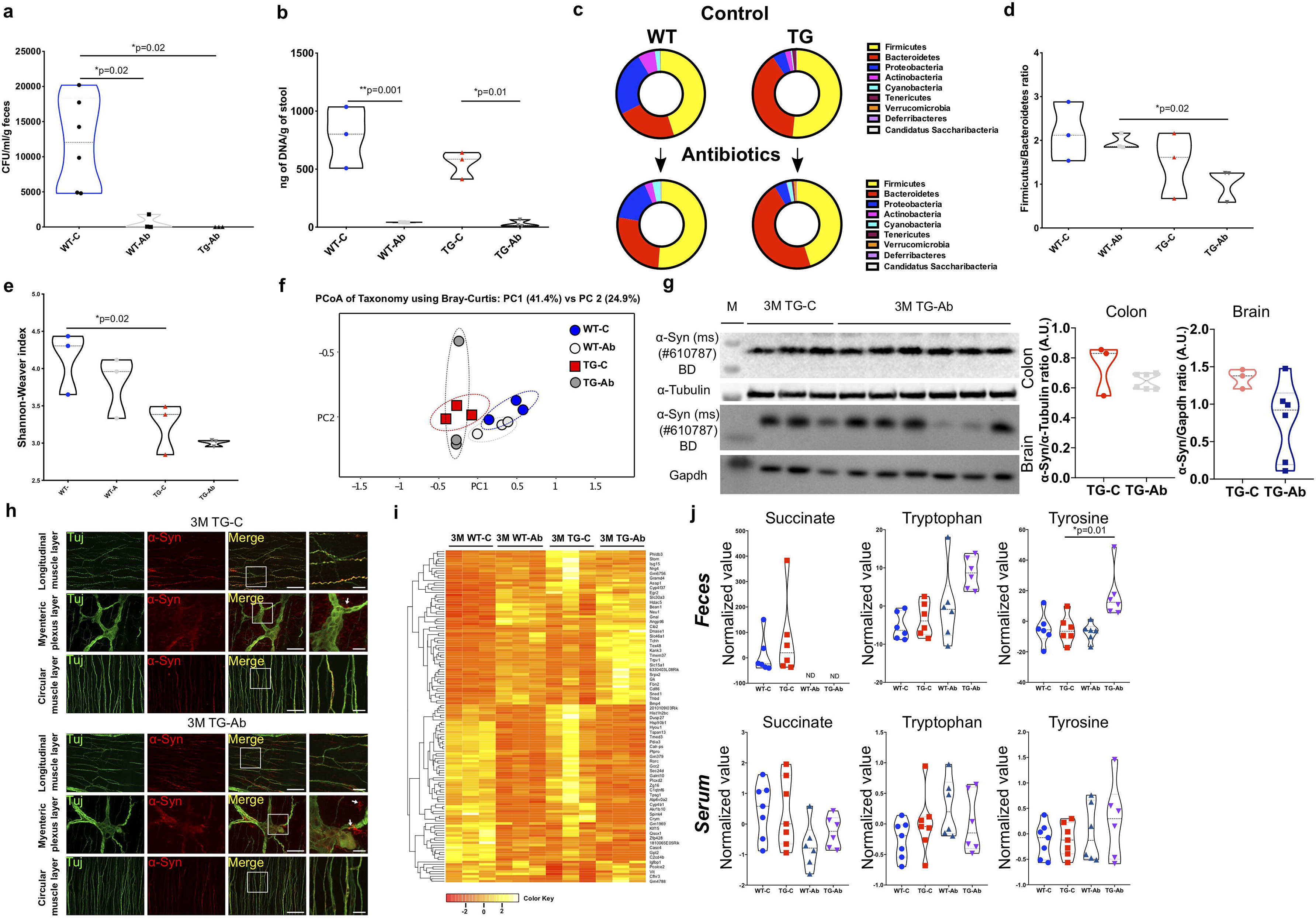
Treatment with broad-spectrum antibiotics leads to reduced gut microbiome load, no change in α-Syn expression and reduced level of succinate metabolites. (a) CFU measurement after broad-spectrum antibiotics treatment in WT and TG rats after equal amount of feces. (b) Amount of total DNA after antibiotics treatment in WT and TG rat faecal samples. (c) Antibiotics affected the bacterial phyla differently for WT and TG rats. (d) F/B ratio was decreased after antibiotic treatment in WT and TG rats respectively, however, no significance difference was observed. (e) Alpha diversity measurement at genera level after antibiotics treatment for WT and TG rats. (f) Beta diversity measurement (genera level) using Bray-Curtis method. Antibiotics treatments affect the clustering of intestinal bacteria. (g) Immunoblot image of α-Syn expression in ENS and olfactory bulb brain region from control and antibiotics treated TG rats and quantification of immunoblots are shown in violin plots. (h) Whole mount staining of muscular layer (longitudinal, circular and myenteric muscle layer) and expression of neuronal (Tuj) and α-Syn expression. (i) Heat-map presenting the RNA-seq data from control and antibiotics treated 3M rats. (j) Succinate, tryptophan and tyrosine levels in the feces and serum from control and antibiotics treated WT and TG (3M) rats. P value significance represents *p≤0.05.

We isolated the *Tunica muscularis* containing the myenteric plexus and found by immunoblotting that antibiotic treatment did not affect the total α-Syn expression in the ENS. However, there was a tendency for reduced α-Syn expression in the brain olfactory bulb region but not reaching to a significant level. (Fig. 7g). Wholemount staining of the *Tunica muscularis* of TG rats from antibiotic treatment and control groups revealed that the expression of α-Syn was present in the myenteric plexus as well as in nerve fibers in the circular and longitudinal muscle layers and that there was no change in expression in response to the antibiotic treatment (Fig. 7h).

Moreover, we performed the RNA-seq analysis of the colon MS layer, we found that several genes were differentially regulated after antibiotics treatment for WT and TG rats respectively (Fig. 7i). IPA analysis suggested that the antibiotics treatment affected 478 genes in TG rats and 521 genes in WT rats (normalised for both the genotypes independently TG-Ab/TG-C and WT-Ab/WT-C). Out of these genes 165 were common in WT and TG. These changes could be caused either by the reduced gut microbiome load or the direct effects of the applied antibiotics (Suppl. Fig. 14). Interestingly, *Dnase1* and *Trpv1* were two of the genes that were upregulated after antibiotic treatment in TG rats. Genes that were upregulated in both WT and TG rats after antibiotic treatments included *Hdac-5*, *CD86* and *Pink1*, whereas *RoRc*, *Spink4*, *Hsp90b1* were downregulated (Suppl. Fig. 14).

Furthermore, we performed ^1^H-NMR metabolomics of antibiotic treated rats and identified 45 faecal and 38 serum metabolites. We found that antibiotic treatment was followed by profound effect on the faecal SCFAs, amino acids (e.g. tyrosine, tryptophan, lysine was reduced in WT rats after antibiotics, whereas no change was observed in TG rats) and sugars (Suppl. Fig 15a & b) while for serum only slight changes could be observed (Suppl. Fig. 15c & d). Most strikingly, multivariate VIP scores analysis of serum samples (Suppl. Fig 14e) identified most strikingly reduced succinate levels (Suppl. Fig. 15f) alongside decreased occurrence of the ketone body 3-Hydroxybutyrate (Suppl. Fig. 15g) in both WT and TG rats after antibiotics treatment. Tyrosine was significantly increased with ageing in TG rat feces after antibiotic treatment, while Tryptophan also had a similar trend but not reaching statistical significance (Fig. 7j). Succinate was totally absent after antibiotic treatment in the feces, thus, suggesting the potential role of succinate production by bacterial metabolism (Fig, 7j). These data signify that microbiome manipulation affects the bacterial load, gene expression and metabolite production.

## Discussion

Recent emerging reports have investigated gut microbiome dysregulation and its role in the pathogenesis of PD^1,17,18,29,49^. Most of the studies propose that PD patients undergo changes in the relative quantity of particular types of bacterial genera and, due to these changes in bacterial load, the metabolism of PD patients is affected^18,50^. Altered bacterial genera/species and their changes to metabolite production may also inadvertently modulate the immune response ultimately, leading to inflammation in various tissues and cellular components of the gut including enteric neurons, glial cells, and immune cells^31^. However, the data from PD patients are always limited in numbers and often have a high variance in terms of age differences, body mass index, exposure to antibiotics, diet, ethnicity etc. To study and understand how the gut microbiome is involved in PD pathology, the above-mentioned factors need to be controlled to have a clear understanding. Moreover, it would be critical to identify patients in the pre-clinical stage who are not experiencing any motor and non-motor symptoms. To address these issues, rodent models (mouse or rats) are useful for identifying and performing functional and mechanistic research on host-microbiome interactions as they allow manipulation of genome, environment and gut microbiome composition^51^. In this study, using the TG PD rat model, we observed that overexpression of α-Syn modifies the gut microbiome, metabolite production and induces inflammation.

Current findings have demonstrated that the intestinal microbiota interacts with the autonomic and central nervous system *via* diverse pathways, including the ENS and Vagal nerve^31^. Studies in PD patients have described that microbial composition in feces and mucosa were significantly different at multiple levels^1,17,49^. The diversity of faecal bacterial communities was not largely different between PD and healthy control subjects^43^. Significant differences in alpha-diversity between PD and healthy control groups were primarily observed at the phylum level and PD disease duration correlated with the majority of taxa, e.g., Bacteroidetes and Proteobacteria being positively correlated and Firmicutes negatively correlated^1^. Additionally, few patient studies reported that Prevotellaceae family or *Prevotella* genus was less abundant in the PD patients^17,18^. We found that *Prevotella* bacterial levels were higher in younger TG rats compared with WT and that at an older age, bacterial levels were lower in TG PD rats, which is consistent with PD patient studies.

Furthermore, our current findings demonstrate that α-Syn expression leads to change in gut microbiota composition and more specifically the *Lactobacillus* and *Alistipes* genera in early young stage, non-symptomatic younger age *versus* older aged, symptomatic TG rats. Abundance of *Lactobacillus* genera is in contrast^17, 22–26^ with many PD patient studies, whilst some are advocating in favour of this study^18,27,28^. *Alistipes* abundance was increased with ageing in TG rats suggesting an increase in inflammation in the colon. Similarly, like rat PD model, *Alistipes* was reported to be increased in PD patients^28^. Reports in other model system such as colon carcinoma and inflammatory bowel disease suggested that increased abundance of *Alistipes* could lead to enhanced inflammatory phenotype in these models^52–54^. It appears that *Alistipes* could be a slow inducer of inflammation in the colon with ageing TG rats, however, further study is warranted. Moreover, our metabolites and immunophenotyping data highlighted enhanced inflammation comes with ageing in TG rats. *Lactobacillus* is considered anti-inflammatory bacteria^55,56^, subsequently there could be a possibility that a low abundance of this bacteria allows a niche for the development of more opportunistic inflammatory bacteria such *Alistipes* and *Desulfovibrio* genera. Additionally, α-Syn overexpression could be key in governing the abundance of *Lactobacillus* in homozygous TG rats. Our maternal microbiome study could explain a decreased abundance of *Lactobacillus* as heterozygous mothers have an equivalent percentage of WT, while homozygous TG rats have less abundant *Lactobacillus*. Thus, the present data are implying that genetics is a major driving force in shaping host-microbiome interactions in our PD rat model. Further, *in vivo* validation with the individual gut bacterial genera/strain is required to support this notion.

A major PD patho-histological hallmark is the occurrence of eosinophilic cytoplasmic neuronal inclusions (Lewy bodies) as well as α-Syn positive neuronal processes called Lewy neurites^4,15,57^. Immunohistochemical data from PD patients before the onset of disease (2-5 years) suggested that α-Syn positive neurons were present in sigmoid colon mucosa^15^. Thus, as suggested earlier, it is possible that inflammation-induced oxidative stress could lead to misfolding of α-Syn in TG rats, and successively α-Syn pathology is spreading to the brain in a prion-like fashion^3,10,11,13,15,58^. It is also conceivable that changes in intestinal bacteria over the course of ageing lead to an increase in intestinal endotoxins, which could enhance the local or systemic inflammation and induce oxidative stress leading to disruption of intestinal barriers or enteric neuroinflammation^30,59^. In line with this, we found that ageing increased gut and systemic inflammation in TG rats and several metabolites such as succinate, lactate, glutamate and 4-hydroxyphenylacetate in the blood serum and feces of aged TG rats (a systemic route of inflammation). It was recently reported that succinate was highly abundant in urine from PD patients compared with healthy controls, furthermore, succinate was correlated with motor score and hence disease severity^60^. Thus, succinate could be used as a prognostic marker for PD pathology. Additionally, the blood serum levels of other anti-inflammatory metabolites such as phenylacetate and tyrosine/tryptophan were decreased in ageing TG rats. Our data suggest that with ageing, an inflammatory environment is indeed be prominent. The inflammatory niche and dysregulated bacterial composition may accelerate the accumulation or aggregation of phosphorylated α-Syn in the myenteric plexuses of TG rats. This is in keeping with previously described results in the brain which suggested that aged TG rats had higher accumulation of α-Syn^3,36^.

Several chronic autoimmune intestinal diseases such as inflammatory bowel disease, celiac disease, type 1 diabetes, multiple sclerosis, and systemic lupus erythematosus are linked with increased intestinal permeability also referred to as “leaky gut”^61^. Thus, gut leakiness in patients with a genetic susceptibility to PD may be a pivotal early step promoting a pro-inflammatory/oxidative environment contributing to the initiation and/or progression of the PD process^30^. In line with this, our Ussing chamber data highlighted that ageing TG rats tended to have more intestinal permeability and significantly less salt entry into intestinal cells than age-matched WT control rats. Thus, we postulate that a lower uptake to Na^+^ ions from the stool could have deleterious effects on *Lactobacillus.* This is in agreement with a recent study, as increased salt intake reduced the *Lactobacillus* in mice model^62^. Furthermore, serum sodium levels are inversely associated with dyskinesia in Parkinson’s disease patients as lower levels of serum sodium are more likely to have dyskinesia^63^. Nevertheless, further understanding the physiology of ion channels and the gut microbiome would be helpful to understand gut-related onset of PD pathology.

Recently several inflammatory proteins such as calprotectin was found to be upregulated in the feces of PD patients^47,48^. The other inflammatory proteins such MCP1 was also correlated with PD progression^64^ as well as in Parkinson’s disease susceptible murine model DJ-1^33^. Our recent data in PD murine model suggested that several inflammatory proteins were also be present in the feces of TG rats^44^. Similarly, we also found that in the rat model with ageing increases levels of calprotectin levels in the feces and serum. Our RNA-seq data pointed that several inflammatory genes such as NFAT regulation of immune response, neuroinflammation and Th1 immune response pathways were upregulated with ageing and reduction in PD-1, PD-L-1 immunomodulatory pathways, antioxidant pathways and LXR/RXR activation in TG rats. Further, these data were validated using immunophenotyping and found that activated CD4^+^ T cells had increased expression of IFN-γ in TG rats. IFN-γ producing T cells involved in induction of inflammation in the brain and activate the inflammatory microglial cells in the *SNpc* region, thus, in turn leads to neuronal cell death and nitrated α-Syn^65^. Most importantly PD-1 inhibitors involved in neurological toxicities in patient with non-small-cell lung cancers^66^ as well as melanoma patient^67^. Thus, reduction in PD-L-1 pathways could also participate in neurodegeneration including PD as our RNA-seq data from colon tissues suggested that PD-1, PD-L-1 immunomodulatory pathways were drastically reduced in TG rats. However, further validation studies are warranted to understand these pathways in PD pathology.

Our microbiome manipulation study by use of broad-spectrum antibiotics again highlights the importance of the gut microbiome and, in particular its ability to modify metabolite production which in theory could be directly involved in the gut inflammation and, thus PD pathology. The antibiotics treatment completely abolished the SCFAs, reduced the succinate production, and increased the production of the amino acids; tyrosine/tryptophan. Tyrosine is the precursor of dopamine synthesis and tryptophan in serotonin production respectively^68^. Accordingly, it could be conceivable that gut bacteria could be regulating the succinate, tyrosine and tryptophan production and could influence neurodegenerative processes by regulating dopamine and serotonin production. Furthermore, it could also be plausible that gut microbiome manipulation by short-term antibiotics treatment could help to modify metabolite production due to dysregulated gut microbiome in PD patients which might improve PD symptoms. However, further detailed investigations are required before it could be implemented for PD patients.

In summary, overall, our TG rat model was able to detect changes in the gut microbiota, intestinal permeability, α-Syn aggregation in parallel with ageing. These characteristics can similarly be detected in human PD patients. A previous study from our lab also suggested that this model exhibits early alterations in avoidance and novelty-seeking behaviour and late motor decline in TG rats^36^ where most of the bacterial diversity and intestinal permeability are drastically changed. Furthermore, our studies showed that α-Syn aggregation could be detected within the walls of the colon of TG rats before the brain pathology develops.

## Material and Methods

### Animals

The BAC-SNCA TG rats have been described in detail earlier and these animals were kept maintaining on outbred conditions on Sprague Dawley background^36^. The rats were housed 3-4 per cage under a 12 h light-dark cycle with ad libitum access to food and water. All experiments were performed according to the EU Animals Scientific Procedures Act and the German law for the welfare of animals. All procedures were approved (TVA: HG3/18) by the authorities of the state of Baden-Württemberg.

### Antibiotics treatment

WT and TG rats (8-10 weeks age) were treated with a cocktail of antibiotics – amoxicillin/clavulanic acid (4:1; 0.5g/L), Vancomycin (0.5g/L), Neomycin (0.5g/L) Cefuroxime (0.5g/L), Streptomycin (5g/L) and ampicillin (0.5g/L) in drinking water containing (5% sucrose). Rats were kept in IVCs one week before the treatment started and till the end of the experiments. Further during the treatment with antibiotics, rats were monitored twice per day for their health and safety (body weight, water content and visual inspection). All the antibiotics treated rats were healthy, a slight reduction of body weight in the beginning of the treatment was observed, however, after one week there was no difference in body weight. Antibiotics were given in an increasing dose (first 0.25g/L and after 3 days it was switched to 0.5g/L until 2 weeks of treatment). The reason for this was to acclimatize the rats for the antibiotics as of the initial concentration of 0.5g/L, the rats did not drink the water and started to lose weight due to dehydration. Once antibiotics were given in lower concentration (0.25g/L) the rats started to drink properly and after 3 days they were switched to a 0.5g/L antibiotic cocktail.

### Bacterial DNA isolation

Faecal pellets were collected from different aged of animals (one-month - 14 months old rats) depending on the experiments for microbial diversity. To collect the faecal pellet, rats were taken out from their home cages and individually placed into a new cage and waited until rats excreted faecal pellet (normally 2-3 pellets were collected). Within 5 minutes of excretion, faecal pellets were kept in cryotubes and transferred onto dry ice during the collection period and after collection of all samples faecal pellets were kept at −80°C until use.

Bacterial DNA was isolated using the QIAamp FAST DNA stool Mini Kit (#51604, Qiagen, Germany) as described by manufacture’s recommendation. In brief, approximately 140-180 mg of faecal pellets was taken, crushed and kept in 2ml tubes for bacterial DNA isolation (pathogen detection) and samples were kept on ice until all the feces were measured. After measurement of all the feces, 1ml of InhibitEX buffer was added to each faecal containing tube. Tubes were vortexed continuously until the faecal sample is thoroughly homogenized and samples were kept at 70°C for 5 minutes and vortexed for 15 seconds after the incubation. Samples were further centrifuged for full speed for 1 minute to pellet faecal particles obtained from the previous step. 200μl buffer AL was added and samples were vortexed for 15 seconds. This mixture was incubated at 70°C for 10 minutes. After incubation, add 200μl of 100% ethanol was added to the lysate and mixed by vortexing. All the lysate was loaded onto QIAamp spin column and centrifuged at full speed for 1 minute. Each column then was washed with 500μl buffer AW1, AW2 and DNA was eluted in 100ul of ATE buffer and DNA was stored at −20°C.

### 16S rRNA sequencing and data analysis

To compare the diversity and composition of gut microbial community, we isolated bacterial DNA from the feces of the WT and TG rats of the respective age (1M-14M) groups cohorts and performed PCR of the 16S ribosomal RNA gene (variable region V3 and V4). The PCR products were sequenced using next generation sequencing. Sequenced amplicons were trimmed and analysed using bioinformatics tools.

For 16S rRNA gene amplification, 12.5ng of DNA was amplified using 0.2μM of both forward (TCGTCGGCAGCGTCAGATGTGTATAAGAGACAGCCTACGGGNGGVWGCAG) and reverse (GTCTCGTGGGCTCGGAGATGTGTATAAGAGACAGGACTACHVGGGTATCTAATCC; both from Metabion) and KAPA HiFi HotStart Ready Mix (#KK2601, KAPABiosystems). PCR was performed using a first denaturation of 95°C for 3 minutes, followed by 25 cycles of amplification at 95°C for 30 seconds, 55°C for 30 seconds and 72°C for 30 seconds, final elongation at 72°C for 5 minutes and the amplified DNA was stored at 4°C. DNA electrophoresis of samples was used to validate the amplicon specificity.

Samples were then subjected to Agencourt AMPure XP PCR purification system (Beckman Coulter) utilizing Agencourt’s solid-phase paramagnetic bead technology for high-throughput purification of PCR amplicons. Agencourt AMPure XP utilizes an optimized buffer to selectively bind PCR amplicons 100bp and larger to paramagnetic beads. Excess primers, nucleotides, salts, and enzymes can be removed using a simple washing procedure. The resulting purified PCR product is essentially free of contaminants. Further, PCR amplicons were indexed using Nextera XT DNA Library Prep Kit and KAPA HiFi HotStart ReadyMix. PCR was performed using first denaturation of 95°C for 3 minutes, followed by 8 cycles of amplification at 95°C for 30 seconds, 55°C for 30 seconds and 72°C for 30 seconds, final elongation at 72°C for 5 minutes. Samples were purified and then validated using BioAnalyzer (Bioanalyzer DNA1000, Agilent) and 4nM of each library pooled using unique induces before sequencing on a MiSeq (Illumina) and paired 300 bp reads.

The obtained sequence reads were sorted by the unique barcodes and sequences of the barcode, linker, and primers were removed from the original sequencing reads using Fastx toolkit. Sequence reads were aligned using MALT (MEGAN alignment tool) for SILVA in semi-global mode and with percent identity threshold of 90%. Further analysis and visualization were performed using MEGAN-CE as described earlier^44^.

### Shotgun sequencing

Isolated DNA was quantified using Qubit (Thermofisher) and presented comparable optical density quality ratio, concentration 26 – 75 ng/μl and fragment size distribution. Samples were normalized to 0.5 ng/μl in Tris (10mM) and an input of 1 ng was processed following Illumina’s Nextera library preparation protocol. DNA was tagmentated for 5 minutes at 55°C. The resulting DNA was then amplified and indexed using 12 PCR enrichment cycles and a dual combination of barcode primers. After amplification, DNA libraries clean-up was performed using AMPure XP bead purification and were resuspended in resuspension buffer. Resulting libraries presented similar molarity 11 – 25 nmol and were pooled based on their molarity in hybridization buffer and denatured following the Nextera XT protocol. Libraries were sequenced using the Illumina NextSeq HighOutput using 2x 150 bp paired end sequencing, providing 34 – 41 million clusters per sample.

### Measurement of Na^+^ Current and gut permeability using an ‘Ussing Chamber’

ENaC activity was estimated from the amiloride-sensitive potential difference and current across the rat colonic epithelium as described in mouse epithelium layer^69^. After removing the outer serosal and the muscular layer under a microscope, tissues were mounted onto a custom-made ‘mini-Ussing chamber’ with an opening diameter of 0.99 mm and an opening area of 0.00769 cm^2^. The serosal and luminal perfusate contained (in mM): 145 NaCl, 1 MgCl2, 2.6 Ca-gluconate, 0.4 KH2PO4, 1.6 K2HPO4, 5 glucose. To assess ENaC induced current, 50 μM amiloride (Sigma, in DMSO) was added to the luminal perfusate. ENaC activity was estimated from the effect of ENaC blocker amiloride (50 μM) on the transepithelial potential difference and current, which reflects the Na^+^ reabsorption from the colon epithelial layer. Transepithelial potential difference (Vte) was determined continuously and transepithelial resistance (Rte) estimated from the voltage deflections (∆Vte) elicited by imposing rectangular test currents of 1 μA and 1.2 s duration at a rate of 8/min. Rte was calculated according to Ohm’s law.

### Preparation of the colon for histological analysis

The colon samples prepared on ice were fixed for 24 h in 4% PFA and then stored at 4°C in 0.4% PFA. Colon samples were prepared from rats deeply anesthetized by CO_2_ inhalation. Fixed colon samples were then alcohol-dehydrated and embedded in paraffin. Samples were embedded in paraffin blocks using a tissue embedding station and stored at room temperature. Paraffin blocks containing brains were cooled on ice block and cut in 7 μm thick sections using a microtome. Sections were placed in 45°C water bath for flattering, collected on a glass slide, dried in an incubator at 50°C for 1 h and stored at room temperature.

### Immunohistochemistry of the colon section

Sections were deparaffinized in xylene and rehydrated in decreasing ethanol. Optionally, a digestion in 50 μg/ml of PK (315836, Roche) was performed in PK-buffer (10 mM Tris, 100 mM NaCl and 0.1% NP-40) at 37°C for 10–60 minutes to stain insoluble proteins. Heat antigen retrieval was performed for 15 min using a microwave set at 1000 W and using 10 mM sodium citrate (pH 6.0) and sodium citrate was refilled every 5 min to avoid the slides to dry. Sections were then washed 3 times 3 min in Tris buffer saline (TBS; Tris 1M, NaCl 3M and pH 7.5) and incubated 20 min in 0.3% hydrogen peroxide to block endogenous peroxidases. After washing the slides 3 times 3 min in TBS, non-specific binding was blocked for 40 min by treating sections with 5% normal serum of the secondary antibody host animal (goat serum: S-1000, Vector; rabbit serum: S-5000, Vector; donkey serum: 017-000-001, Dianova) diluted in TBS and supplemented with 0.3% Triton X-100 to permeabilize cell membranes. After washing the slides 3 times 3 min with TBS, sections were incubated overnight with the primary antibody diluted in TBS containing 1% normal serum. The next day, slides were washed 3 times 3 min with TBS+0.025% Triton X-100 and incubated for 1 h with a biotin conjugated secondary antibody (mouse: BA-9200, rabbit: BA-1000, rat: BA-4000; Vector) diluted 1:250 in PBS containing 1% normal serum. Sections were washed 3 times 3 min with TBS+0.025% Triton X-100 and then incubated with an avidin-biotin enhancer complex coupled with peroxidase (ABC Elite; Vector) at room temperature for 1 h. After washing 3 times 3 min with TBS+0.025% Triton X-100, sections were detected using 3,3′-diaminobenzidine (DAB; Sigma) providing a brown precipitate when oxidized by the peroxidase linked to the secondary antibody. Reaction was stopped by washing slides in distilled water after 3 min. For microscopy, slides were mounted using a cover slipping media, dried overnight, and stored at room temperature.

### Western blotting

The colons for the mass spectrometry or Immunoblotting analysis were collected from freshly euthanized rats and cleaned with PBS to remove faecal material and snap-frozen in liquid nitrogen and samples were stored at −80°C freezer until use. In certain experiments, myenteric plexus and epithelial layer were also isolated and samples were stored at −80°C until use.

Proteins were lysed from the colon tissues in 10 volume of RIPA buffer (50 mM Tris, 150 mM NaCl, 1.0% NP-40, 0.5% sodium deoxycholate, 0.1% SDS, pH 8.0) supplemented with protease inhibitor (Complete; Roche Diagnostics) in order to perform gel electrophoresis. Brain tissues were disrupted 30 sec using a homogenizer (T10 ultra turrax; VWR) in ice. After the homogenization, samples were incubated for 30 min at 4°C and spun for 20 min at 12000xg. Proteins lysate supernatants were supplemented with 10% glycerol before long storage at −80°C.

Protein concentration was determined using BCA method (BCA Protein Assay Kit, 23225, Life Technology) or Bradford assay. Samples were prepared by diluting protein lysates in PAGE buffer (0.2 M glycine, 25 mM Tris, 1% SDS), followed by a denaturation at 95°C for 10 min in loading buffer (80 mM Tris, 2% SDS, 5% 2-mercaptoethanol, 10% glycerol, 0.005% bromophenol blue, pH 6.8) and a short centrifugation 30 sec at 400×g. Proteins were separated by electrophoresis using 12% SDS-PAGE gel. Gels containing proteins were washed for 5 minutes in transfer buffer (0.2 M glycine, 25 mM Tris, 10–20% methanol) and transferred to membranes equilibrated in transfer buffer. Transfer was performed for 1h 30 minutes at 80 V at 4°C on PVDF membranes. Immunoblot were washed 5 min in TBS buffer and blocked using 5% non-fat milk (Slim Fast) in TBS. Membranes were then washed twice 5 min in Tris buffer saline containing 0.01% Triton X-100 (TBST) and incubated with the primary antibody over night at 4°C (human and mouse a-Syn: 610786 BD Biosciences). After incubation with the first antibody, membranes were washed four times (5 min each) with TBST. Membranes were then incubated for 75 min with the secondary antibody coupled to horseradish peroxidase (GE Healthcare). After four washing steps with TBST (5 min each), bands were visualized using the enhanced chemiluminescence method (ECL+; GE Healthcare). Light signal was detected using LI-COR Odyssey and were quantified using Odyssey software.

### ELISA

To measure the calprotectin/MRP 8/14 from the fecal samples, S100A8/S100A9 ELISA kit (#K6936, Immundiagnostik AG) was used according to the manufacture’s guidelines. First faecal samples were measured (weight 50 mg) and dissolved in 500 ul of extraction buffer supplied by the kit, mixed by vortexing and then centrifuged for 10 min at 3000xg. Supernatant was taken and transferred to a new microcentrifuge tube and 100 μl of sample was used for measuring the protein. The data were analyzed using 4 parameters algorithm and the concentration of calprotectin was normalized with feces weight and data are presented in ng/g.

### RNA-sequencing from the colon tissues

Total RNA was extracted using QIAsymphony RNA kit (#931636, Qiagen). Briefly, 10 to 20 mg of frozen tissue was dissociated using 400 μl of RLT Plus in a 2mL extraction tube containing a 5 mm diameter beads (Qiagen) and agitated at 30 HZ for twice 2 minutes in the TissueLyser II (Qiagen). RNA isolation was performed on the QIAsymphony (Qiagen) following the platform Standard protocol. Elution was performed using 50 μl of RNase-free water.

RNA quality was assessed with an Agilent 2100 Bioanalyzer and the Agilent RNA 6000 Nano kit (Agilent). 3’RNA-sequencing was performed using 100 ng of total RNA and the QuantSeq 3’ mRNA-Seq (#015.24; Lexogen, Austria). Libraries were sequenced on the NextSeq500 using the Mid Output v2.5 150 Cycles (#20024904, Illumina) with a depth of >2 millions reads each.

FASTQ were generated using fastp (v0.20.0) and RNA-seq data quality was assessed to identify potential issues with sequencing cycles, low average quality, adaptor contamination, or repetitive sequences from PCR amplification. Reads were aligned using STAR (v2.7.2a) against a custom-built genome composed of the Ensembl Mus musculus grcm38. Alignment quality was analyzed using MappingQC (v1.8) and visually inspected in the Integrative Genome Viewer (v2.4.19). Normalized read counts for all genes were obtained using edgeR (v3.26.6). Transcripts covered with less than 4 count-per-million in at least 1 sample were excluded from the analysis leaving >14,000 genes for determining differential expression in each of the pair-wise comparisons between experimental groups.

RNA sequencing data was further subjected to Ingenuity pathway analysis (IPA) to identify the genes involved in different canonical pathways as well as with reactome online tool.

### Flow cytometry

Plasma cells were first washed with PBS at 600xg for 5 minutes at 4 °C. Supernatant was discarded, and cells were first stained with surface staining antibodies (CD4, CD8, CD25, CD44) then were fixed with Foxp3 fixation/permeabilization buffer (eBioscience, Germany) for intracellular staining and incubated for 30 minutes. After incubation, cells were washed with 1x permeabilization buffer, exposed to added intracellular monoclonal antibodies (Foxp3, IL-17 and IFN-g in different channel) and incubated for an additional 45 minutes. Cells were washed again with permeabilization buffer and PBS was added to acquire the cells on a flow cytometer (BSFortessa™ from Becton Dickinson; Heidelberg, Germany).

### Whole-mount procedure and staining

Tunica muscularis strips were fixed with 4 % (w/v) phosphate buffered p-formaldehyde (PFA; Sigma-Aldrich, St. Louis, MO, USA) for 20 min and rinsed three times with phosphate buffered saline (PBS). Antigen retrieval was carried out in citrate buffer (0.01 mol/l, pH 6.0, Roth, Karlsruhe, Germany) by heating the samples in a microwave (600 W for 5 min) and successively allowed the samples cool overnight. To prevent unspecific binding of antibodies, samples were blocked for two days at room temperature with PBS containing 4 % (v/v) goat serum (Biochrom, Berlin, Germany), 0.1% (v/v) bovine serum albumin (BSA; Roth), 0.3 % (v/v) Triton® X-100 (Roth), and 0.1% (wt/v) NaN_3_ (Merck, Darmstadt, Germany) on a shaker. Afterwards, samples were incubated with primary antibody diluted in PBS with 0.1% (v/v) BSA, 0.3% (v/v) Triton^®^ X-100, and 0.1% (wt/v) NaN_3_ for three days at room temperature. Samples were then rinsed in PBS with 0.1% (wt/v) NaN_3_ three times, with the last washing step over night. Then secondary antibodies were diluted in PBS with 0.1% (v/v) BSA, 0.3% (v/v) Triton® X-100, and 0.1% (wt/v) NaN_3_ and incubated overnight. Samples were then washed three times in PBS, immersed in 80% (v/v) glycerol (Sigma-Aldrich) in PBS for refractive index matching, and placed in a glass bottom petri dish for imaging.

The following primary antibodies were used for immunocytochemistry: mouse anti-α-Synuclein (1:500, BD, Franklin Lakes, NJ, USA), rabbit anti-β-III tubulin (TUJ, 1:400, BioLegend, San Diego, CA, USA). Primary antibodies were detected using the fluorescent secondary antibodies goat anti-rabbit Alexa 488 (1:400, Invitrogen, Carlsbad, CA, USA) and goat anti-mouse Alexa 546 (1:400, Invitrogen). Nuclear staining was carried out with DAPI solution (200 ng/ml, Roth). A Zeiss Axio Imager.Z1 fluorescence microscope with integrated Apotome module (Zeiss, Jena, Germany) was used for microscopic evaluation and structured illumination imaging. Images were acquired using Zeiss Zen blue software (Zeiss).

### Feces and serum sample preparation for the metabolite detection using ^1^H-NMR

For metabolite extraction and separation from lipids that disturb the ^1^H-NMR spectra of polar metabolites, a standard 2-phase extraction protocol was used. Hereby, 50 mg of deep-frozen feces sample were transferred into 2 ml adaptive focused acoustics (AFA) glass tubes (Covaris Inc, Woburn, USA) and mixed with 400 μl of ultrapure methanol and 800 μL of Methyl ter-butyl ether (MTBE; solvent grade). The mixture was manually dispersed with a disposable plastic spatula, then vortexed and transferred to a focused ultrasonicator (Covaris E220evolution, Woburn, USA). Feces metabolites were extracted with a 5 min lasting ultrasonication program in a degassed water bath at 7° C. After extraction, the metabolite suspension was separated into a polar and lipid phase by adding 400 μl of ultrapure water. In order to remove any remaining solids from the samples, the glass tubes were centrifuged for 5 min at 4000xg. 700 μl of polar upper phase was then transferred to a fresh 1.5 ml eppendorf tube. The polar phase was subject to a 2nd centrifugation step for 5 min at 12,000xg and 600 μl of the supernatant were transferred to a new 1.5 ml eppendorf tube and evaporated to dryness over night with a vacuum concentrator (Eppendorf Speedvac).

We prepared the serum samples for NMR analysis using following method. Briefly, 45 μl of serum and 90 μl of 100% methanol was used and then mixed with the Covaris 130 microTUBE glass vials for metabolites extractions (no phase separation was used). This procedure was done for 25 minutes (5 minutes/samples) in one cycle. Once the mixing was done, 135 μl of sample mixture supernatant was collected in a fresh 500 μl tube and subjected to centrifuge for 30 minutes at 12,000xg. After centrifugation, 120 μl of supernatant was taken in a fresh 1.5 ml Eppendorf tube and subjected to speed vacuum for the overnight and dried pellets was dissolved in 50 μl NMR buffer (recipe+STDs) was added and briefly sonicated to mix the buffer and metabolite pellets. Mixed solution 45 μl of the supernatant were transferred with gel loading pipette tips into 1.7 mm NMR tubes (Bruker BioSpin, Karlsruhe, Germany) and a 96 well rack placed into the cooled (4° C) NMR autosampler.

For the feces samples, we used a slightly different protocol for NMR analysis. The dried faecal pellets (after overnight speedvac) were resuspended with 60 μl of deuterated phosphate buffer (200 mM K2HPO4, 200 μM NaN3, pH 7.4) containing 1 mM of the internal standard TSP (trimethylsilylpropanoic acid). For a maximum dissolution, the plastic tubes were quickly sonicated and then centrifuged 5 min at 14,000xg. 50 μl of the supernatant were transferred with gel loading pipette tips into 1.7 mm NMR tubes (Bruker BioSpin, Karlsruhe, Germany) and a 96 well rack placed into the cooled (4° C) NMR autosampler.

Spectra were recorded on a 600 MHz ultra-shielded NMR spectrometer (Avance III, Bruker BioSpin GmbH) equipped with a triple resonance (^1^H, ^13^C, ^31^P) 1.7 mm room temperature probe at 298 K. For optimum water suppression and shim adjustment a quick simple ZG experiment was performed followed by a 1hour lasting CPMG (Carr-Purcell-Meiboom-Gill) experiment in order to suppress residual background signals from macromolecules such as bilirubin (time domain = 64k points, sweep width = 20 ppm, 512 scans). The recorded free induction decays (FIDs) were fourier-transformed and spectra properly phase- and baseline corrected (Bruker Topspin 3.5.6). The hereby created files were processed with ChenomX NMR Suite 8.3 for metabolite annotation and quantification. For statistical analysis, different grouped metabolite concentration tables were created and imported with MetaboAnalyst 4.0. In detail, we investigated both the effect of ageing (between 3 and 14 months) as well as genotype (WT and TG) on the metabolic phenotypes in serum and feces.

### Statistical analysis

MEGAN-CE (version 6.14.2, built 23 Jan 2019) and MicrobiomeAnalyst were used for data acquisition and analysis. GraphPad and Inkscape were used for the final figure preparation. One-way ANOVA, Two-way ANOVA (post-hoc Sidak’s test) or Student’s t-test was used for statistical analysis using GraphPad (version 8) wherever it was appropriate and described in the figure legend. Data shown in violin plots which represents the median and quartiles as the box-and-whisker plots and it also display a smoothed frequency distribution. The p value (≤0.05) considered significant. Metabolite concentrations from ^1^H-NMR analysis were exported as comma separated value spreadsheet file to MetaboAnalyst, normalized with PQN (probabilistic quantile normalization) to account for dilution effects and pareto scaled for making metabolites within one sample comparable. A combined fold change (FC > 1.2) and Student’s t-test analysis (p < 0.05) was used for the volcano and box plots. Furthermore, multivariate heat maps, PLS-DA plots and VIP scores were generated for direct comparison of the genotypes at 3M and >12M in feces and serum, respectively. The effect of antibiotics treatment on serum and feces metabolite profiles was investigated by heat maps, sPLS-DA and VIP scores.

## Supporting information

Suppl. Fig-YSINGH

## Declarations

### Ethics statement and approval

All the experiments were performed according to the EU Animals Scientific Procedures Act (2010/63/EU) and the German law for the welfare of the animals. All the procedure and methods were approved by the local government authorities (Regierungspräsidium, Tübingen; TVA HG3/18) of the state of Baden-Württemberg, Germany.

### Consent for publication

No patients or human data was used in this study. All authors read the manuscript and approved to be co-authors on the manuscript and have substantial contribution in the manuscript.

### Availability of data and material

The datasets used and/or analysed during the current study are available from the corresponding authors on a reasonable request. RNA-seq data are available from GEO accession no. xxx.

### Competing interests

The authors declare that they have no competing interests.

### Funding

This research project is an EU Joint Programme - Neurodegenerative Disease Research (JPND) (JPCOFUND_FP-829-047 aSynProtect) and is supported through the funding organization Deutschland, Bundesministerium für Bildung und Forschung (BMBF, FKZ). This research was also supported by the Deutsche Forschungsgemeinschaft, DFG through the funding of the NGS Competent Centre Tübingen (NCCT-DFG). Funders have no role in the study design and data analysis.

## Acknowledgements

We thank Prof Bernd Pichler for allowing to have access to high-field ^1^H-NMR spectroscopy for metabolomics studies. We acknowledge Prof. Jeroen Raes, VIB-KU Leuven Center for Microbiology, Rega Institute, Leuven, Netherlands for helpful discussion with microbiome data and calprotectin assay development. We would like to acknowledge the support from FACS core facility and NGS Competence Centre Tübingen (NCCT). We also acknowledge the helpful discussion with aSynProtect team members (Jia-Yi Li, Tiago Outeiro, Olaf Riess, Pekka Kallunki, Daniel Otzen, Richard Wade-Martins, Jeroen Raes, Hilal Lashuel). We acknowledge support by Deutsche Forschungsgemeinschaft (DFG) and Open Access Publishing Fund of University of Tübingen.

## Author’s contribution

YS: Study design, performed the research and managed the overall project, involved in entire study, analyzed the data, made the figures, and wrote the manuscript.

CT: Performed the metabolites study, data analysis, made the figures, wrote the manuscript

JR, HAL, K-HS: performed the characterization of a-Syn from the ENS of the rat colon, expression of a-Syn in whole-mount staining and analysed the data.

AD, JA: Helped with 16S rRNA data analysis bioinformatics meta data analysis and made the figures.

MSS, FL: performed the Ussing chamber experiments and analysed the data.

PHN: Performed the whole-mount staining procedure and microscopy for antibiotic treatment experiments.

IF, MA, NW: performed the Immunoblotting experiments from colon and brain samples and data analysis.

MDS, DO: Study design for microbiome sequencing and data analysis.

NC, OR: Provided tools for the study, original microbiome study design, data analyses, discussion, edited the manuscript and funding acquisition.

## Suppl. Figures

Suppl. Fig. 1 The gut microbiome dynamics with ageing, increased *Lactobacillus* and decreased *Alistipes* bacterial genera in the TG rats. (a) 16S rRNA gene amplicon sequencing was performed on samples from 1, 2, 2.5, 3, 6 and 14 M age group WT and TG respectively. Total number of sequences read obtained from WT and TG samples were similar and a significant difference was not observed. (b) The alpha diversity within each group (WT and TG rat) was estimated using Shannon-Weaver index based on phylum level classifications. No significant difference was noticed in between WT and TG rats at any age group. Data are represented in Box and Whisker plots. (c) Age dynamics of phyla Firmicutes, Bacteroidetes, Actinobacteria, Proteobacteria and Verrucomicrobia. Each phylum was significantly different at particular age as shown in the figure with significant level. (d) The Shannon-Weaver index and Chao 1 (alpha diversity) at genus level at particular age. Significant change is shown in asterisk for a defined age group. (e) The dynamic representation of *Prevotella, Alloprevotella, Parabacteroides, Alistipes Lactobacillus, Turicibacter, Ruminococcus, Desulfovibrio* and *Bifidobacterium* with ageing in WT and TG respectively. A significant difference is shown for a particular age group. P value significance represents *p≤0.05, **p≤0.01 and ***p≤0.001.

Suppl. Fig. 2 Clustering of bacterial phyla for different age group WT and TG rats. Bacterial clustering figure showed change in the bacterial abundance with ageing.

Suppl. Fig. 3 Genetic makeup of animal could be an important factor for microbiome composition development. (a) A study was carried out from 4M old heterozygous females (total no of female (F) 6; 2 females/cage (C); colour coded) were kept in 3 different cages and when females were pregnant then male were separated. Faecal samples were collected from the females after pups were born. After 3 weeks of age, pups were separated from females and genotyped and kept in WT and homozygous TG groups in 5 different cages (C1-5) as explained in the fig. Faecal samples were sequenced by 16S rRNA sequencing and analysed by MEGAN-CE software. Clustering analysis at species level suggested that WT and homozygous TG rats’ clusters differently with each other as well as with mother’s (Heterozygous) microbiome. (b) The percentage abundance of *Lactobacillus* genus and species in two different facilities at the same age group in two different cohorts. P value significance represents *p≤0.05.

Suppl. Fig. 4 Similar presence of *Lactobacillus* genus in two different facilities. (a) At 3M age species comparison study showed that two different facilities had similar abundance of *Lactobacillus* species. (b) The percentage abundance of *Lactobacillus* genus and species in two different facilities at the same age group in two different cohorts.

Suppl. Fig. 5 Coprophagy is one of route and method for microbial manipulation in PD rat models (a) Schematic diagram for bacterial manipulation by cage transplant. (b) Total no of sequence reads. (c) Hollow pie charts represent the bacterial composition at phylum level before and after faecal transplant. (d) F/B ratio before and after faecal transplant. (e) Pie charts show the bacterial composition before and after faecal transplant in WT and TG PD rat model. (f) Bacterial composition in individual WT and TG rats before and after transplant and represented as heat-map.

Suppl. Fig. 6 The gut microbiome dynamics with ageing, increased *Lactobacillus* and decreased *Alistipes* bacterial genera in TG rats. (a) Total no of sequence reads after shotgun sequencing obtained from WT an TG samples were similar. (b) Rarefaction plots for all samples. (c) Clustering of bacterial at species level from WT and TG rats (>12M). (d) Significantly changed bacterial species in WT and TG rats.

Suppl. Fig. 7 The metabolic profiles of 3M feces (a) and serum (c) show no clear separation of WT and TG rats in the PLS-DA scores plots and also heat maps show no clear clusters. By contrast, after >12M TG and WT samples could be clearly separated in the PLS-DA scores plots in both feces (b) and serum (d) analysis. The most prominent metabolites in the VIP scores analysis for feces samples were succinate (high in TG for 3M) and glutamate (high in WT for >12M). For serum, 3M rats showed high glucose levels in TG and high lactate in WT, while at >12M TG rats showed highly increased lactate and succinate levels.

Suppl. Fig. 8 The serum metabolome analysis of WT rats (a) revealed significant (p < 0.05) metabolite fold changes (FC > 1.2) during ageing. At >12M of lifetime, WT serum were composed with increased levels of formate, isoleucine, valine, taurine, creatine, glutamine and lysine and decreased values of betaine, N,N-dimethylglycine, glutamate, sn-glycero-3-phosphocholine, citrate and 3-hydroxybutyrate. Succinate was increased at 3 months of age, however above the significance threshold. Results from the same spectra with multivariate PLS-DA VIP scores analysis (c) provided similar results for WT while TG rats showed mainly an in increase in lactate and succinate. A comparison of the feces metabolome during aging showed both for WT and TG a decrease in succinate levels (b) for which a correlation analysis was performed.

Suppl. Fig. 9 Fragmented α-Syn and expression in ENS from the colon of >12M TG rats. (a) Estimation of fragmentation in α-Syn in from the colon ENS. Two major truncated fragments were found in ENS as shown in α-Syn immunoblot image. (b) Expression of α-Syn in the colon muscular layer.

Suppl. Fig. 10 Accumulation of α-Syn in TG rat colon. Colon tissues were digested with trypsin for 2 hours (1:5 dilution of the original) and performed with human (TG rats) and rat (WT rats) specific antibodies as described in materials and methods. IHC showed the accumulation of α-Syn in both 2M and 18M TG rats, whereas in WT (for both the age group) α-Syn was dissolved by trypsin so it appeared that α-Syn in WT existed in soluble form whereas α-Syn was retained in TG rats suggested the accumulation.

Suppl. Fig. 11 Human α-Syn is truncated in TG rats. (a) The expression of endogenous α-Syn in rat colon and upto 4 different truncated α-Syn can be observed in the colon of the TG rats, but the appearance of these different bands differs between individual and does not seem to be correlated with age. In some TG rats (5M 1 and 18M 1), the expression of α-Syn is similar than in the WT rats but showing truncated fragments of lower molecular weight. No pathologic phosphorylation (S129, Y125, Y39) could be detected by immunoblotting method. (b). DSP treatment increases α-Syn immunodetection in WB. The detection of the truncated α-Syn band is not dependent on boiling the samples, as similar band is also present in non-boiled samples, but its signal is increased in presence of the crosslinker DSP. (c) Using different α-Syn antibodies covering different epitopes of the sequence were used to detect the fragmentation of α-Syn. N-terminal antibodies did not detect the first truncated fragment present in sample 1 and 2-Detect the second truncated fragment present in sample 2 and 3. Detect an extra truncated fragment in sample 1 with a similar molecular weight the second truncated fragment. C-terminal antibodies do not detect the first and second truncated fragment present in sample 1 and 2, but only upto residue 125. SA3400 (117-131) does not detect these truncated forms. The extra truncated fragment in sample 1 is detected by all c-terminal antibodies. 1^st^ truncated band is observed with α-Syn 211 (121-125), 14H2L1 (117-125), LB509(115-122) and FL140 (61-95), however it was not observed with SA3400 (117-131), EP1466Y (N-term), N19 (N-term) and Ab6176 (11-26). The fact that the 1^st^ truncated band was not observed by SA3400 (117-131), however it was observed by all the other C-term antibodies upto 125, suggest that it might be a N-terminal fragment with the truncation located between the position 125 and 131. Further, the truncated fragment was not observed by any of the three N-term antibodies tested, thus further suggesting that the fragment was also truncated at N-terminal part. The 2^nd^ band was observed with N19 (N-term), Ab6176 (11-26), EP1646Y (N-term), 14H2L1 (117-125) and LB509 (115-122) and not observed with SA3400 (117-131). Thus, detected band was observed by all the N-term antibodies but not by the most distant C-term antibody (SA3400) suggesting that it was truncated in the C-terminal part, probably around the position 120. (d) A cartoon is showing possible aberrant alternative splicing variant sites.

Suppl. Fig. 12 RNA-seq analysis from 3M and >12M old WT and TG colon epithelial tissues. (a) Differentially expressed genes at 3M age in WT and TG rats. Total no of differentially expressed genes (153) based on absolute log fold-change of at least 1 (raw p value P=0.05), whereas at >12M age total no of differentially expressed were 1808 based on absolute log fold-change of at least 1 (raw p value P=0.05). There was no statistical significance based on false discovery rate (FDR) for 3M, however at >12M 264 genes were different. (b) Clustering of >12M WT and TG epithelial (Epi) samples. (c,d), IPA analysis of adaptive (Th1 pathway) and innate (TLR pathway) for the >12M samples. (e) Different inflammatory gene expressions

Suppl. Fig. 13 Reactome transcriptome wide overview analysis of 3M and >12M epithelial RNA-seq samples. The figure shows a genome-wide overview of the results of pathway analysis (a) 3M and (b) >12M. Reactome pathways are arranged in a hierarchy. The center of each of the circular “bursts” is the root of the one top-level pathway. Each step away from the center presents the next level lower in the pathway hierarchy. The colour code denotes over-representation of a particular pathway in the input dataset. Light grey signifies pathways which are not significantly over-presented. At 3M programmed cell death pathway genes were over-presented whereas at >12 M programmed cell death pathways gene were downregulated. Further, at >12M M there were an over-representation of immune system genes and less representation of programmed cell death genes.

Suppl. Fig. 14 Antibiotics treatment increase TRPV1 and Dnase1 in the colon. (a) Clustering of the samples in the control (WT and TG) and antibiotics treated groups. (b) Genotype analysis in control and antibiotics treated colon samples (epi). Total 165 genes were found to be commonly regulated by antibiotics. More differential gene expression in WT compared with TG rats. (c) Expression of TRPV1, Dnase1 in WT and TG colon.

Suppl. Fig. 15 Metabolites changes in the feces and serum after antibiotics treatment at 3M age. (a) Antibiotics treatment results in a clear separation of feces samples in the (a) PLS-DA analysis (b) and heat map. (c) While the effect on the corresponding serum scores plots and (d) heat map is rather small. (e,f,g) Multivariate VIP scores analysis identified decreased succinate in both TG and WT control samples after antibiotics treatment alongside the ketone body 3-Hydroxybutyrate. Box and whisker plots showed the succinate and 3-Hydroxybutyrate in control and antibiotic treated samples.

## Notes

### Competing Interest Statement

The authors have declared no competing interest.

