## Supplementary material for "Overexpression of human alpha-Synuclein leads to dysregulated microbiome/metabolites with ageing in a rat model of Parkinson disease": Suppl. Fig-YSINGH

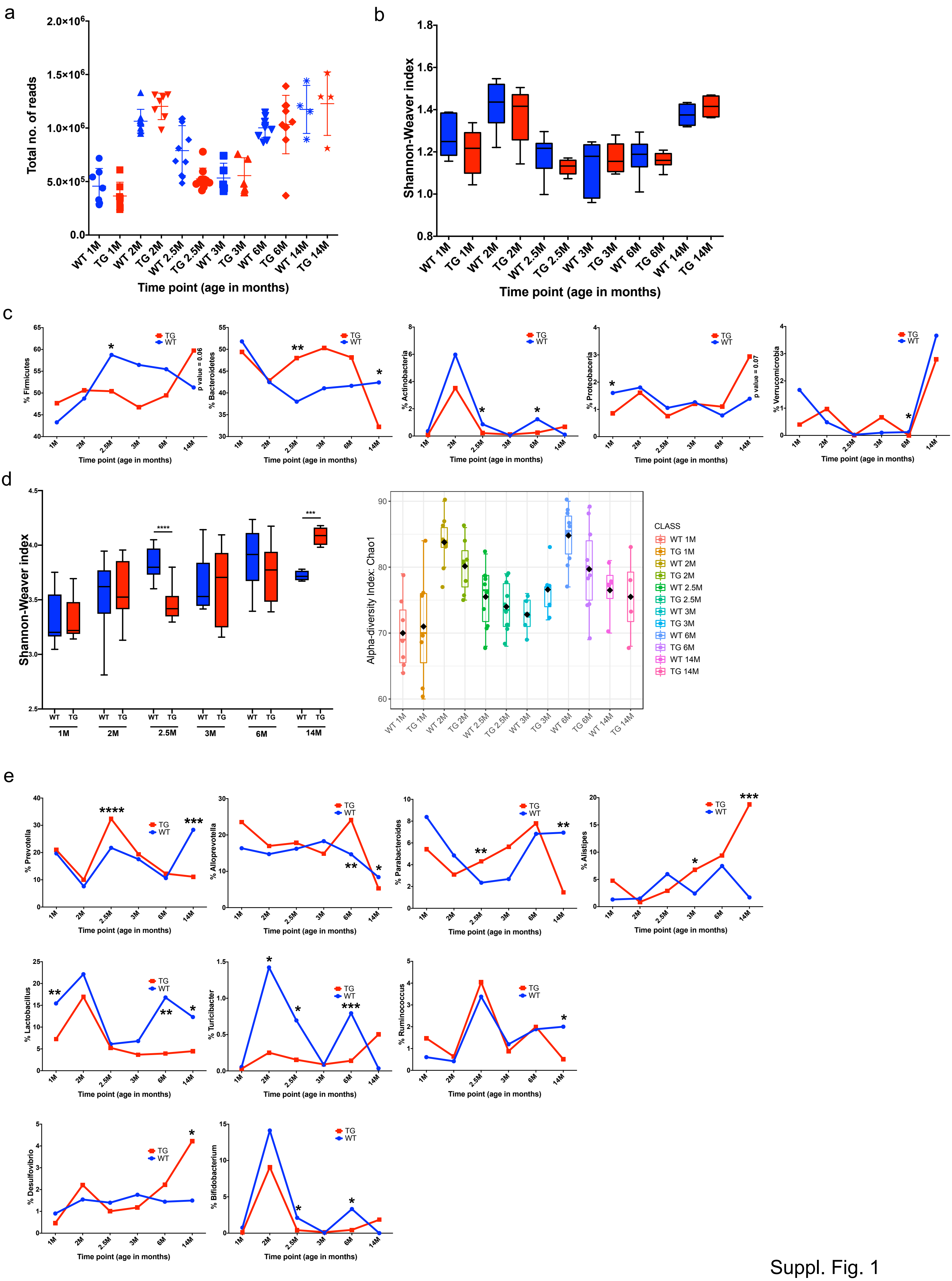
a
Total no. of reads
Time point (age in months)
b
Shannon-Weaver index
Time point (age in months)
c
% Firmicutes
% Bacteroidetes
p value = 0.06
% Actinobacteria
% Proteobacteria
p value = 0.07
% Verrucomicrobia
TG
WT
d
Shannon-Weaver index
Alpha-diversity Index: Chao1
CLASS
e
% Prevotella
% Alloprevotella
% Parabacteroides
% Alistipes
% Lactobacillus
% Turicibacter
% Ruminococcus
% Desulfovibrio
% Bifidobacterium

Suppl. Fig. 1

Microbial dynamics at the genus level in WT and TG rats

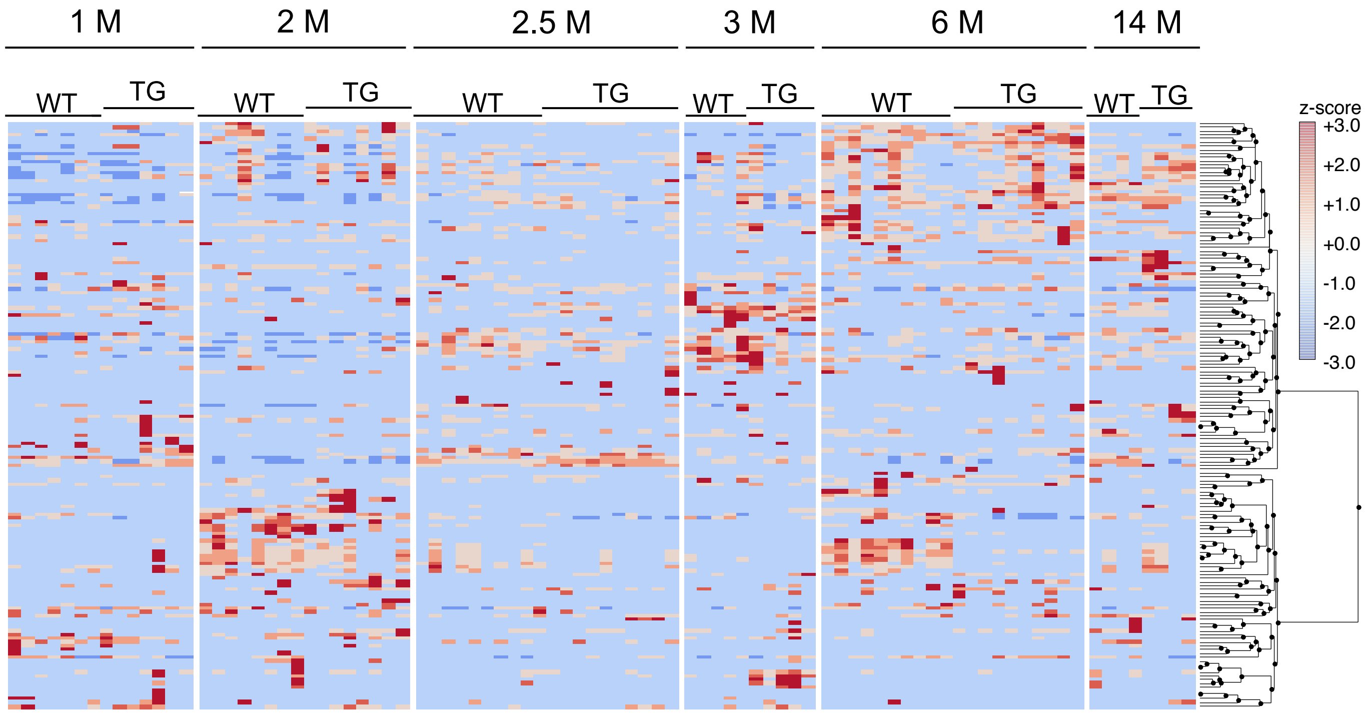

Suppl. Fig. 2

a

Mother to children microbiome study

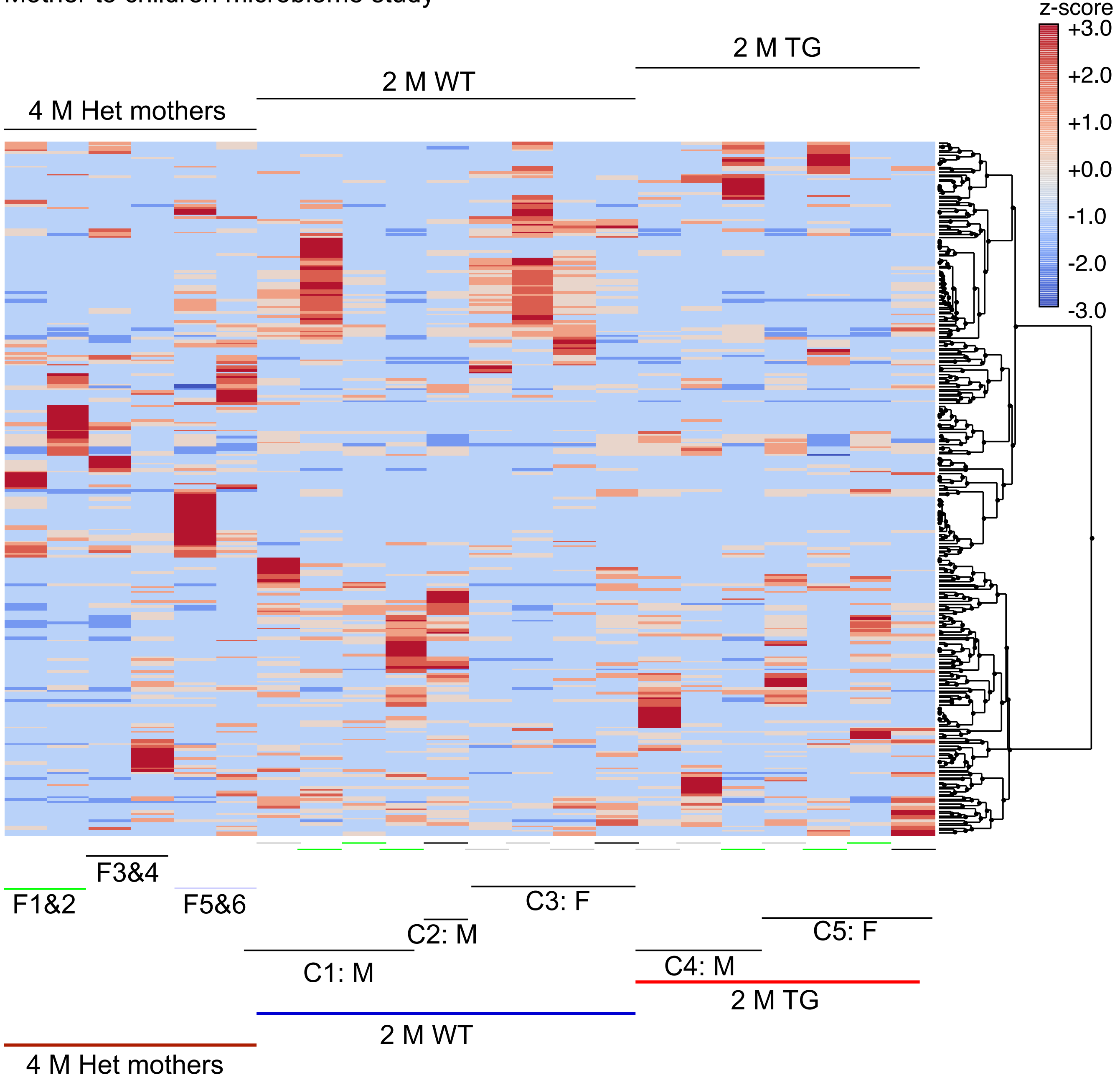

b

| Series: Genus (% abundance) | Het 4M | WT 2M | TG 2M |
|---|---|---|---|
| Lactobacillus | 22.0699983 | 22.5240711 | 17.4533753 |

c

| Series: Species (% abundance) | Het 4M | WT 2M | TG 2M |
|---|---|---|---|
| uncultured Lactobacillus sp. | 0.03805911 | 0.10114158 | 0.03175778 |
| Lactobacillus acidophilus | 0.00632911 | 0.02617774 | 0.00675949 |
| Lactobacillus brevis | 0.01146372 | 0.03944909 | 0.01417334 |
| Lactobacillus buchneri | 0 | 0.00641777 | 0.00102447 |
| Lactobacillus casei | 0.00183167 | 0.0316849 | 0.01058442 |
| Lactobacillus paracasei | 0.01140339 | 0.01952132 | 0.00391501 |
| Lactobacillus coleohominis | 0 | 0.00087215 | 0 |
| Lactobacillus crispatus | 0.0101515 | 0.02857682 | 0.00923365 |
| Lactobacillus delbrueckii | 0.0030155 | 0.02211619 | 0.00234591 |
| Lactobacillus fermentum | 0 | 0.00573385 | 0.00117083 |
| Lactobacillus frumenti | 0 | 0.00352063 | 0 |
| Lactobacillus gallinarum | 0 | 0.00468202 | 0 |
| Lactobacillus helveticus | 0.02348155 | 0.08417008 | 0.02419469 |
| Lactobacillus hilgardii | 0.01508796 | 0 | 0 |
| Lactobacillus iners | 0.03719946 | 0.00604975 | 0 |
| Lactobacillus jensenii | 0.00323888 | 0.00193079 | 0 |
| Lactobacillus johnsonii | 0.25728342 | 0.08022391 | 0.10624953 |
| Lactobacillus kefiranofaciens | 0.00516473 | 0.01771349 | 0.00315086 |
| Lactobacillus kunkeei | 0.00279571 | 0.01066086 | 0.001244 |
| Lactobacillus malefermentans | 0 | 0.00521185 | 0 |
| Lactobacillus oris | 0 | 0.02997847 | 0.00433518 |
| Lactobacillus panis | 0.00269931 | 0.01142564 | 0.00242633 |
| Lactobacillus pentosus | 0 | 0.01971269 | 0.00437977 |
| Lactobacillus plantarum | 0.16481063 | 0.89659491 | 0.29772974 |
| Lactobacillus pontis | 0 | 0.00275847 | 0 |
| Lactobacillus reuteri | 0.01379851 | 0.0269457 | 0.00874326 |
| Lactobacillus sp. AcjLac17 | 0 | 0.00587665 | 0 |
| Lactobacillus sp. Akhmbto4 | 0 | 0.00414891 | 0 |
| Lactobacillus sp. GV6 | 0.00263439 | 0.00194903 | 0 |
| Lactobacillus sp. lab13 | 0 | 0.00095937 | 0 |
| Lactobacillus sp. N3-1-1 | 0.00134965 | 0.00354556 | 0 |

Suppl. Fig. 3

a

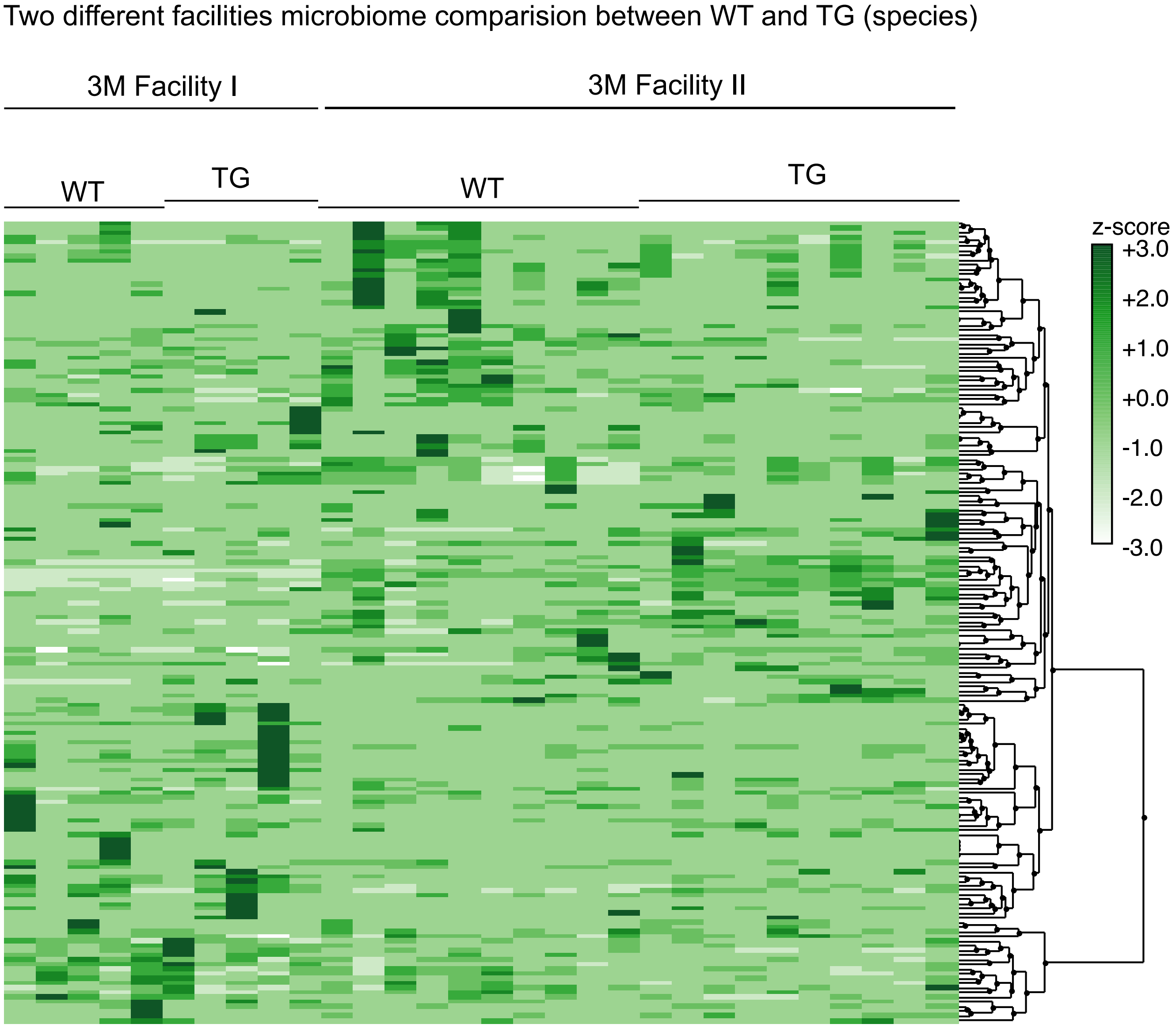

b

| #Series: Genus (% abundance) | WT 3M FI | TG 3M FI | WT 3M FII | TG 3M FII |
|---|---|---|---|---|
| Lactobacillus | 7.13507466 | 3.86075136 | 6.43720752 | 5.42566682 |
| #Series: Species (% abundance) | WT 3M FI | TG 3M FI | WT 3M FII | TG 3M FII |
| Lactobacillus acidophilus | 0.00145057 | 0 | 0.00205119 | 0.00047017 |
| Lactobacillus brevis | 0.00096705 | 0 | 0.00357856 | 0.00218986 |
| Lactobacillus casei | 0.00103482 | 0 | 0 | 0 |
| Lactobacillus crispatus | 0.00352641 | 0.00112983 | 0.00258844 | 0.00180516 |
| Lactobacillus delbrueckii | 0 | 0 | 0.00055984 | 0 |
| Lactobacillus helveticus | 0.00550231 | 0.00169475 | 0.00985339 | 0.00529313 |
| Lactobacillus hilgardii | 0 | 0 | 0.00425296 | 0.00210576 |
| Lactobacillus iners | 0.00224363 | 0 | 0.01329042 | 0.0084205 |
| Lactobacillus johnsonii | 0.05668325 | 0.04552587 | 0.08163448 | 0.06776931 |
| Lactobacillus kefiranofaciens | 0.00096705 | 0 | 0.00174272 | 0 |
| Lactobacillus panis | 0.00096705 | 0 | 0 | 0 |
| Lactobacillus paracasei | 0.00145057 | 0 | 0.00293537 | 0.00051986 |
| Lactobacillus plantarum | 0.06776078 | 0.04501744 | 0.06961048 | 0.04198644 |
| Lactobacillus reuteri | 0 | 0 | 0.00384287 | 0.0025342 |
| Lactobacillus sp. GV6 | 0.00096705 | 0.00112983 | 0.00062571 | 0 |

Suppl. Fig. 4

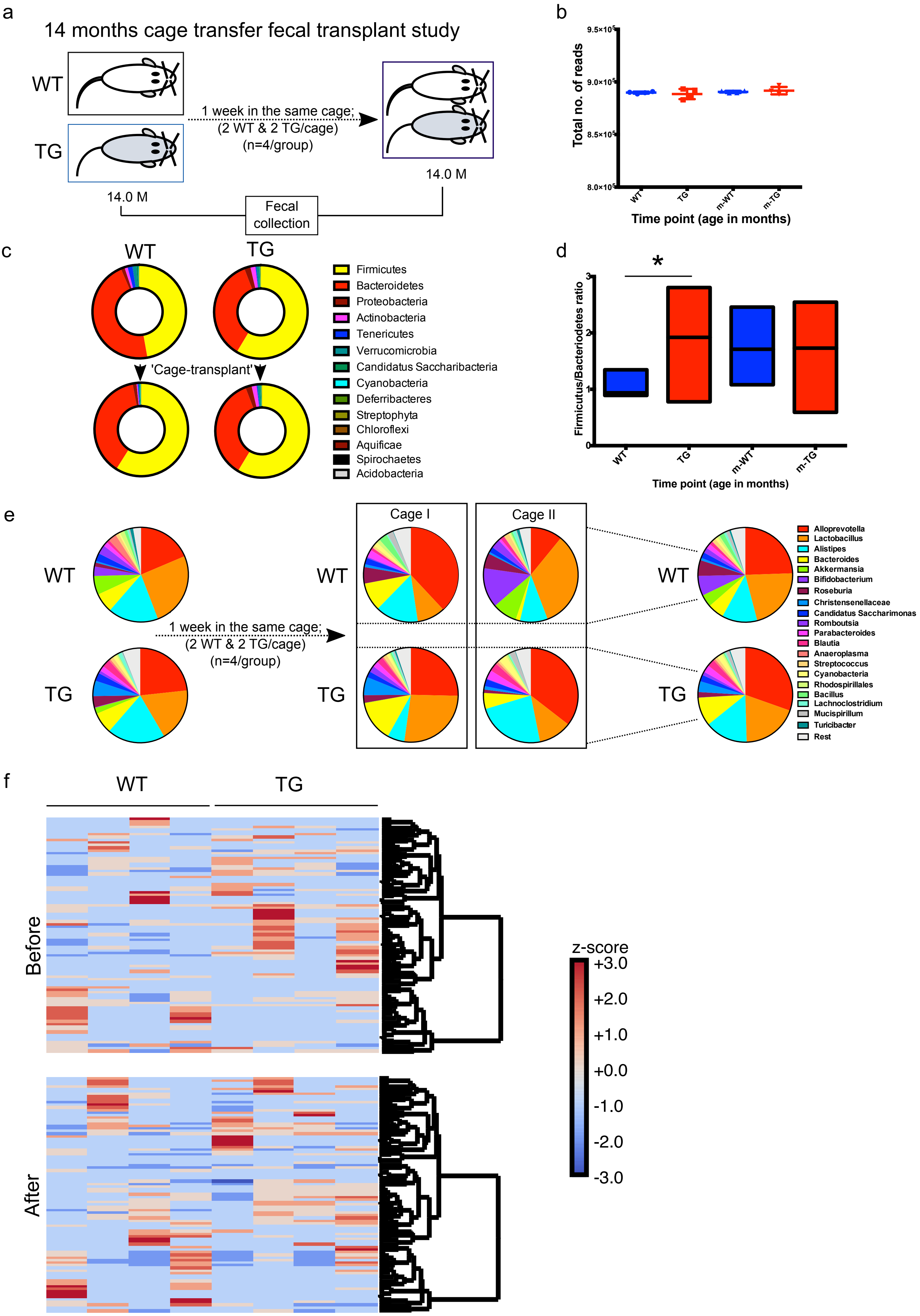
a
14 months cage transfer fecal transplant study
WT
TG
1 week in the same cage;
(2 WT & 2 TG/cage)
(n=4/group)
14.0 M
14.0 M
Fecal collection
b
Total no. of reads
9.5×10^5
9.0×10^5
8.5×10^5
8.0×10^5
WT
TG
m-WT
m-TG
Time point (age in months)
c
WT
TG
'Cage-transplant'
Firmicutes
Bacteroidetes
Proteobacteria
Actinobacteria
Tenericutes
Verrucomicrobia
Candidatus Saccharibacteria
Cyanobacteria
Deferribacteres
Streptophyta
Chloroflexi
Aquificae
Spirochaetes
Acidobacteria
d
*
Firmicutus/Bacteriodetes ratio
WT
TG
m-WT
m-TG
Time point (age in months)
e
WT
TG
1 week in the same cage;
(2 WT & 2 TG/cage)
(n=4/group)
Cage I
Cage II
Alloprevotella
Lactobacillus
Alistipes
Bacteroides
Akkermansia
Bifidobacterium
Roseburia
Christensenellaceae
Candidatus Saccharimonas
Romboutsia
Parabacteroides
Blautia
Anaeroplasma
Streptococcus
Cyanobacteria
Rhodospirillales
Bacillus
Lachnoclostridium
Mucispirillum
Turicibacter
Rest
f
WT
TG
Before
After
z-score
+3.0
+2.0
+1.0
+0.0
-1.0
-2.0
-3.0

Suppl. Fig. 5

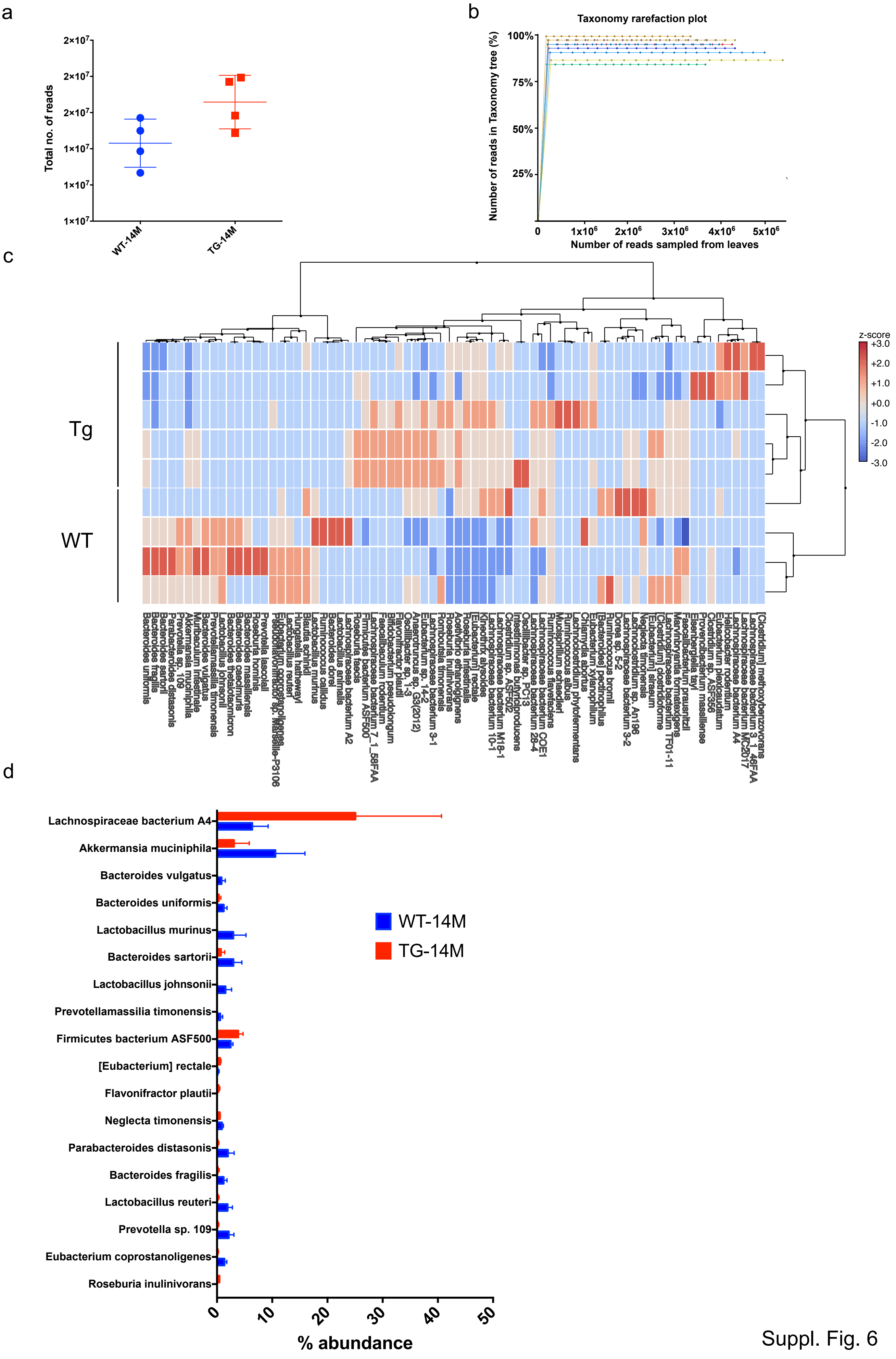

a
b
c
d
Taxonomy rarefaction plot
Number of reads in Taxonomy tree (%)
Number of reads sampled from leaves
Total no. of reads
WT-14M
TG-14M
Tg
WT
z-score
Lachnospiraceae bacterium A4
Akkermansia muciniphila
Bacteroides vulgatus
Bacteroides uniformis
Lactobacillus murinus
Bacteroides sartorii
Lactobacillus johnsonii
Prevotellamassilia timonensis
Firmicutes bacterium ASF500
[Eubacterium] rectale
Flavonifractor plautii
Neglecta timonensis
Parabacteroides distasonis
Bacteroides fragilis
Lactobacillus reuteri
Prevotella sp. 109
Eubacterium coprostanoligenes
Roseburia inulinivorans
% abundance

Suppl. Fig. 6

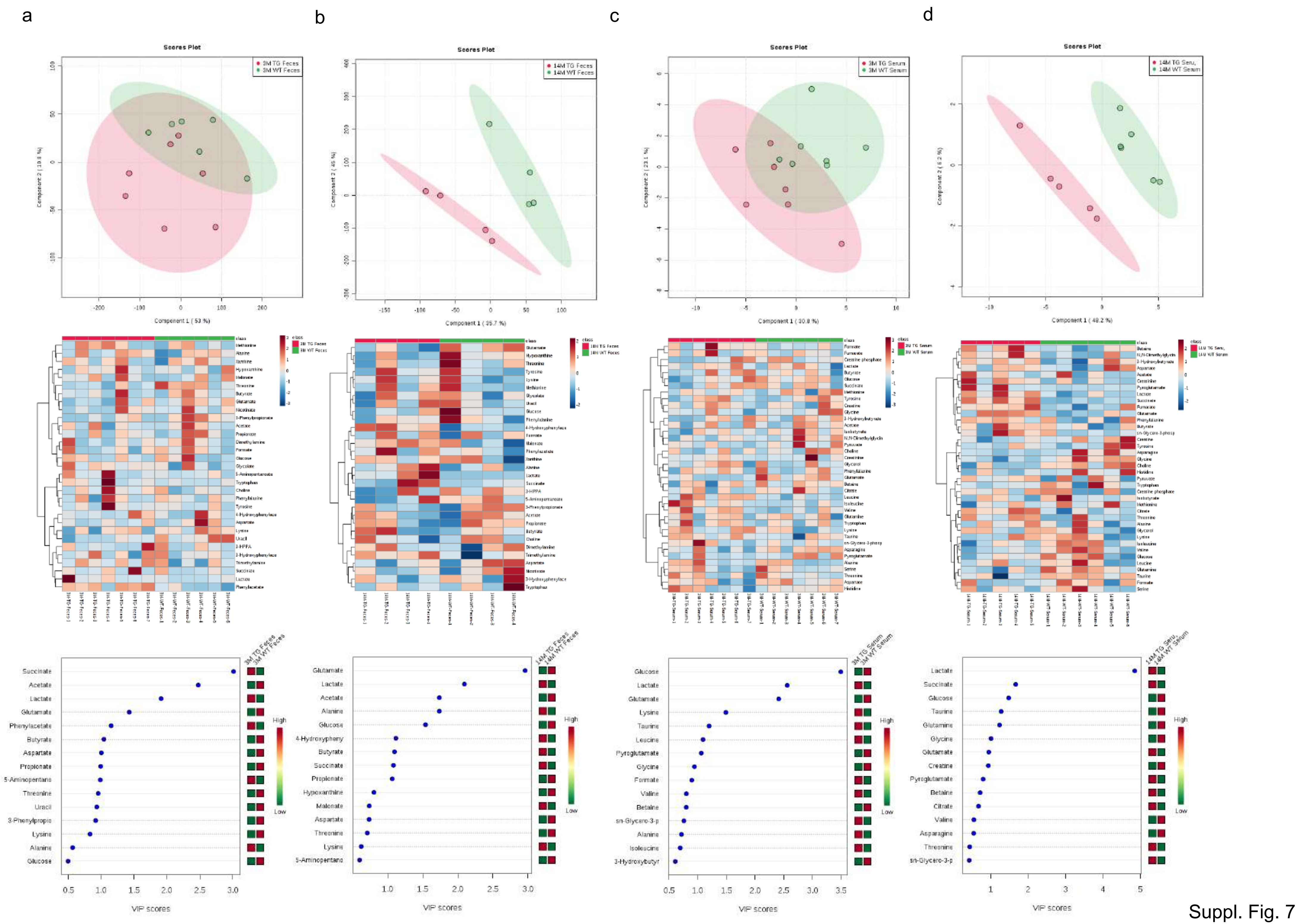
a
b
c
d
Scores Plot
3M TG Feces
3M WT Feces
14M TG Feces
14M WT Feces
3M TG Serum
3M WT Serum
14M TG Seru,
14M WT Serum
VIP scores
High
Low

Suppl. Fig. 7

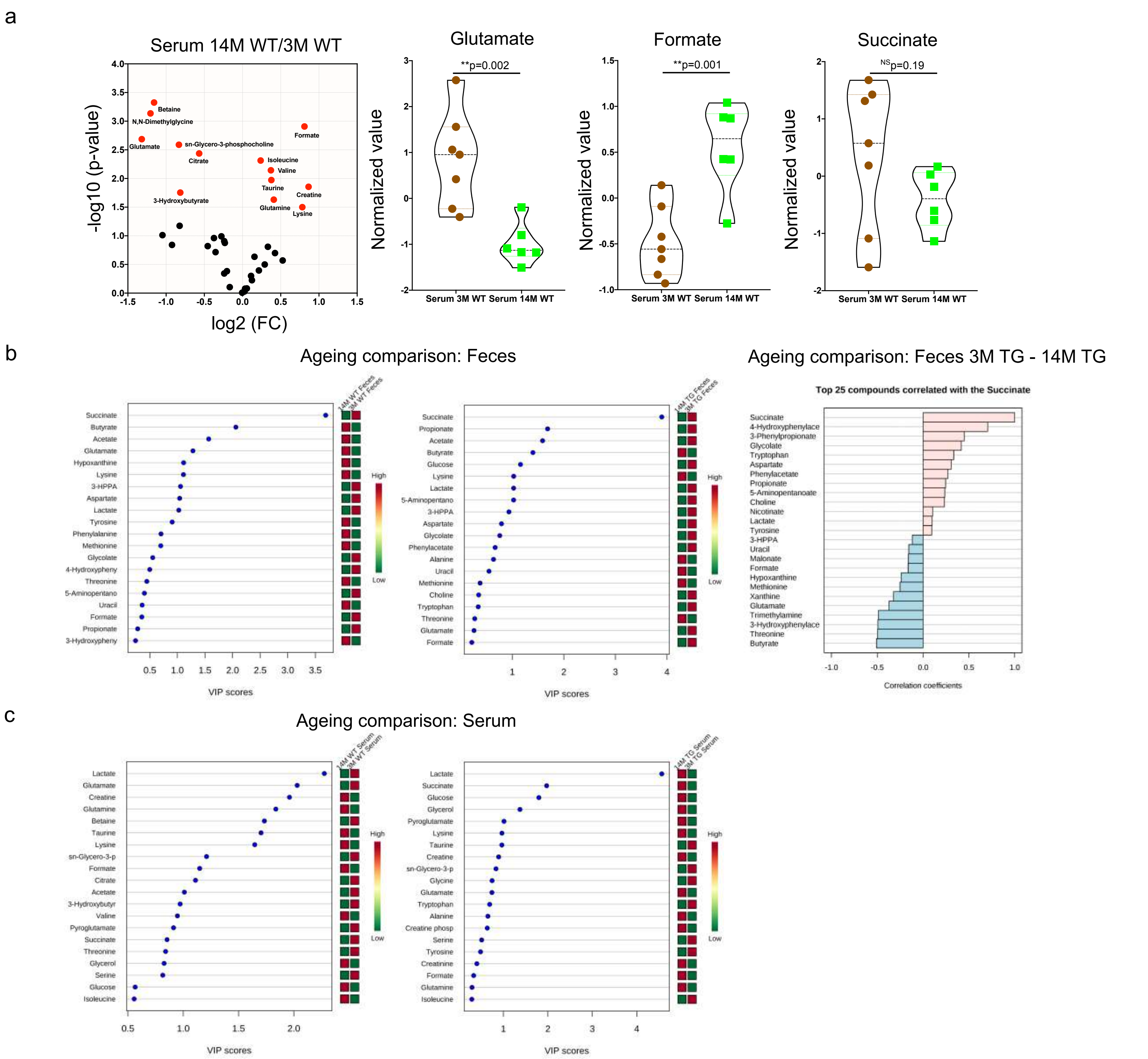
a
Serum 14M WT/3M WT
-log10 (p-value)
log2 (FC)
Betaine
N,N-Dimethylglycine
Glutamate
sn-Glycero-3-phosphocholine
Citrate
Formate
Isoleucine
Valine
Taurine
Creatine
3-Hydroxybutyrate
Glutamine
Lysine
Glutamate
**p=0.002
Formate
**p=0.001
Succinate
NSp=0.19
Normalized value
Serum 3M WT Serum 14M WT
b
Ageing comparison: Feces
14M WT Feces
3M WT Feces
Succinate
Butyrate
Acetate
Glutamate
Hypoxanthine
Lysine
3-HPPA
Aspartate
Lactate
Tyrosine
Phenylalanine
Methionine
Glycolate
4-Hydroxypheny
Threonine
5-Aminopentano
Uracil
Formate
Propionate
3-Hydroxypheny
VIP scores
High
Low
14M TG Feces
3M TG Feces
Succinate
Propionate
Acetate
Butyrate
Glucose
Lysine
Lactate
5-Aminopentano
3-HPPA
Aspartate
Glycolate
Phenylacetate
Alanine
Uracil
Methionine
Choline
Tryptophan
Threonine
Glutamate
Formate
VIP scores
Ageing comparison: Feces 3M TG - 14M TG
Top 25 compounds correlated with the Succinate
Succinate
4-Hydroxyphenylace
3-Phenylpropionate
Glycolate
Tryptophan
Aspartate
Phenylacetate
Propionate
5-Aminopentanoate
Choline
Nicotinate
Lactate
Tyrosine
3-HPPA
Uracil
Malonate
Formate
Hypoxanthine
Methionine
Xanthine
Glutamate
Trimethylamine
3-Hydroxyphenylace
Threonine
Butyrate
Correlation coefficients
c
Ageing comparison: Serum
14M WT Serum
3M WT Serum
Lactate
Glutamate
Creatine
Glutamine
Betaine
Taurine
Lysine
sn-Glycero-3-p
Formate
Citrate
Acetate
3-Hydroxybutyr
Valine
Pyroglutamate
Succinate
Threonine
Glycerol
Serine
Glucose
Isoleucine
VIP scores
14M TG Serum
3M TG Serum
Lactate
Succinate
Glucose
Glycerol
Pyroglutamate
Lysine
Taurine
Creatine
sn-Glycero-3-p
Glycine
Glutamate
Tryptophan
Alanine
Creatine phosp
Serine
Tyrosine
Creatinine
Formate
Glutamine
Isoleucine
VIP scores

Suppl. Fig. 8

a

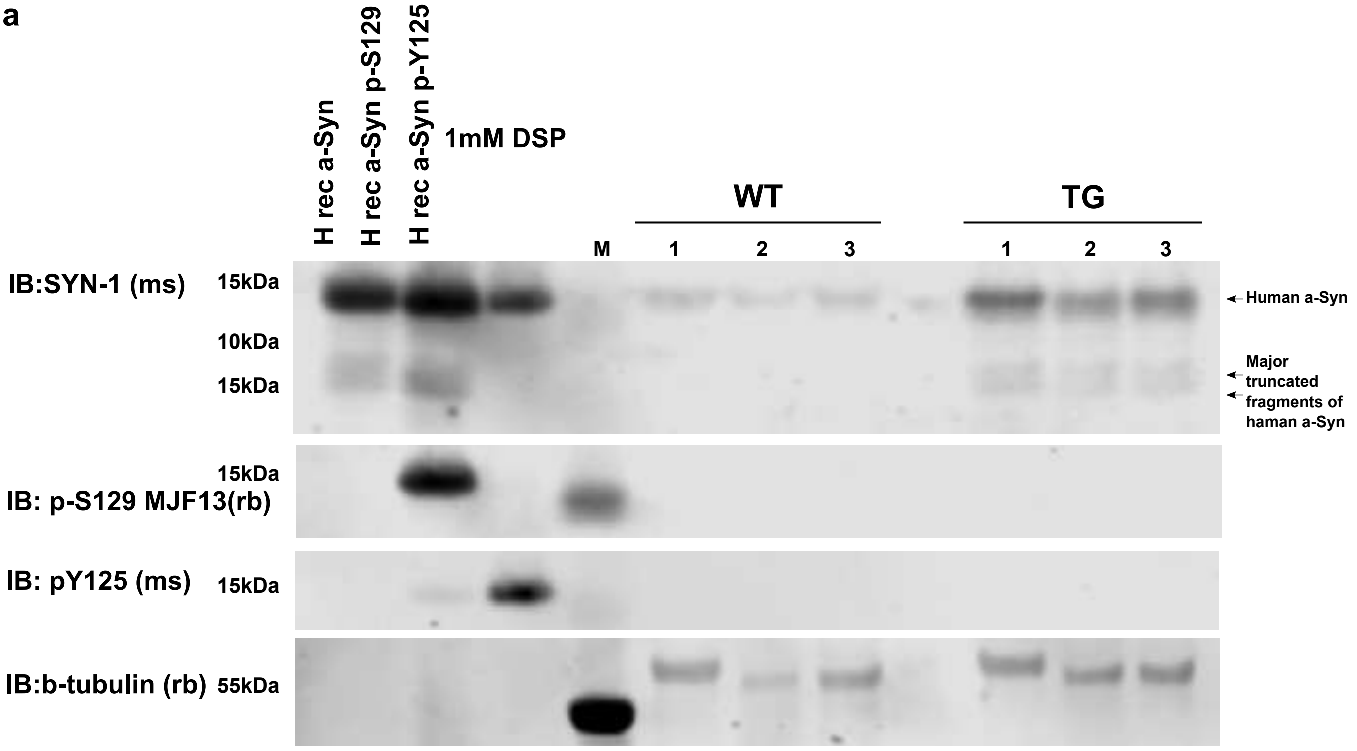

b

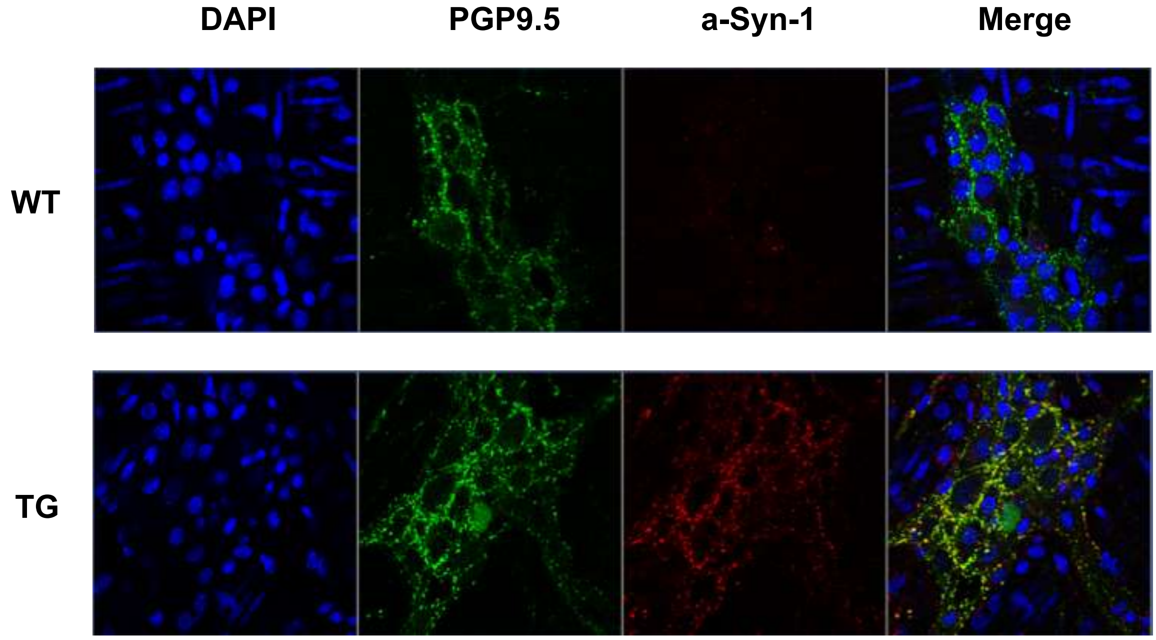

Suppl. Fig. 9

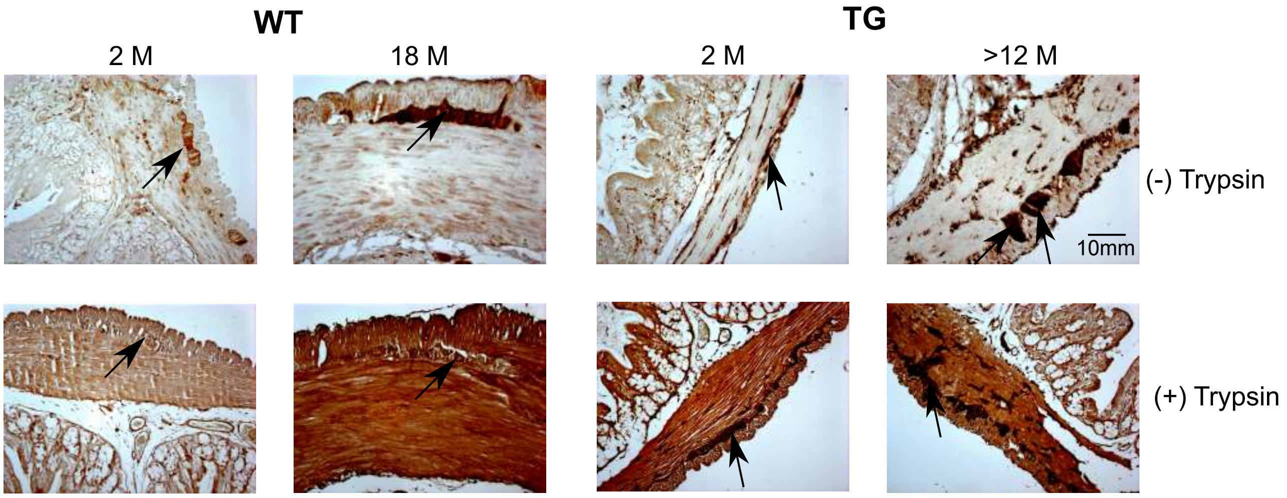

Suppl. Fig. 10

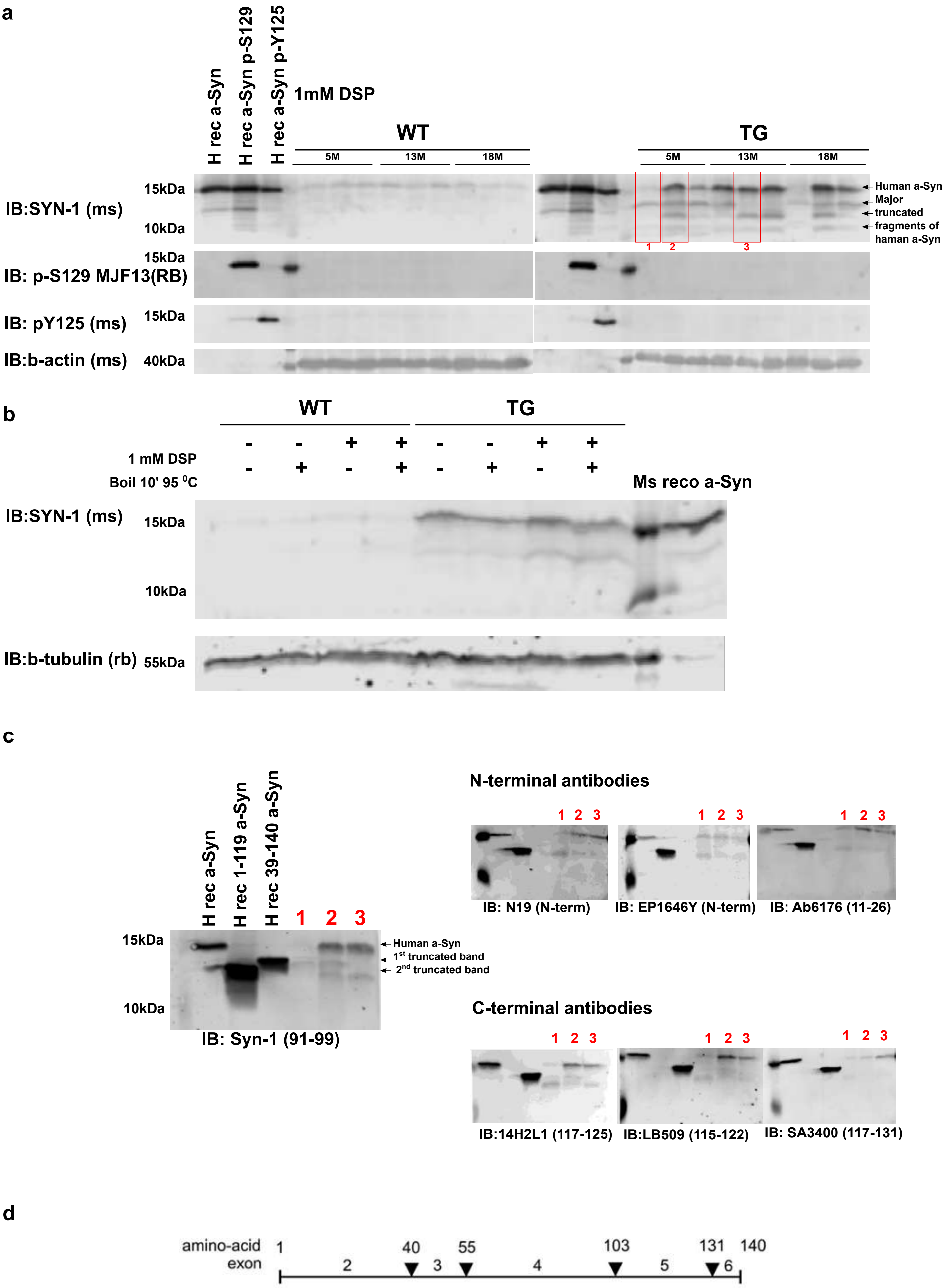
a
H rec a-Syn
H rec a-Syn p-S129
H rec a-Syn p-Y125
1mM DSP
WT
TG
5M
13M
18M
IB:SYN-1 (ms)
15kDa
10kDa
Human a-Syn
Major truncated fragments of haman a-Syn
1
2
3
IB: p-S129 MJF13(RB)
IB: pY125 (ms)
IB:b-actin (ms)
40kDa
b
1 mM DSP
Boil 10' 95 °C
Ms reco a-Syn
IB:b-tubulin (rb)
55kDa
c
H rec 1-119 a-Syn
H rec 39-140 a-Syn
1st truncated band
2nd truncated band
IB: Syn-1 (91-99)
N-terminal antibodies
IB: N19 (N-term)
IB: EP1646Y (N-term)
IB: Ab6176 (11-26)
C-terminal antibodies
IB:14H2L1 (117-125)
IB:LB509 (115-122)
IB: SA3400 (117-131)
d
amino-acid
exon
1
40
55
103
131
140
2
3
4
5
6

Suppl. Fig. 11

**a**

| Contrast | *FC | #FDR |
|---|---|---|
| 3M (TG/WT) | 153 | 0 |
| 14M (TG/WT) | 1808 | 264 |

***FC=** absolute log fold change of at least 1, i.e. double or half expression strength

**#FDR=** statistical significance (FDR/adjusted p-value) of at most 0.05

**b**

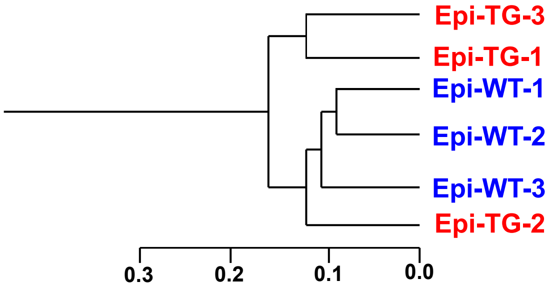

**c**

Th1 signaling pathways: Adaptive immunity

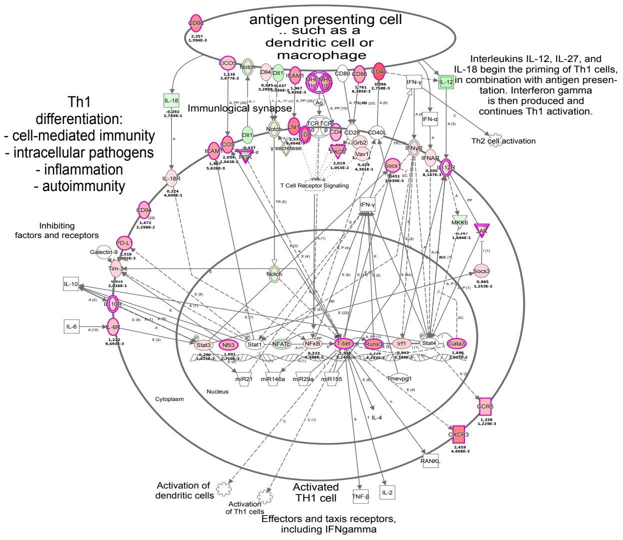

**d**

Toll like receptor signaling pathways: Innate immunity

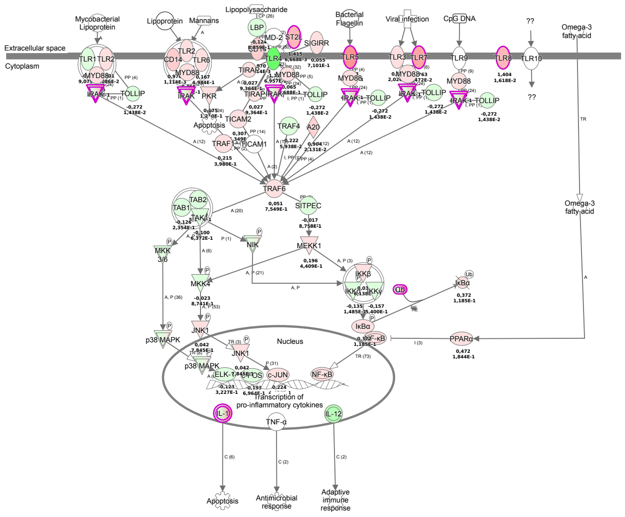

Suppl. Fig. 12

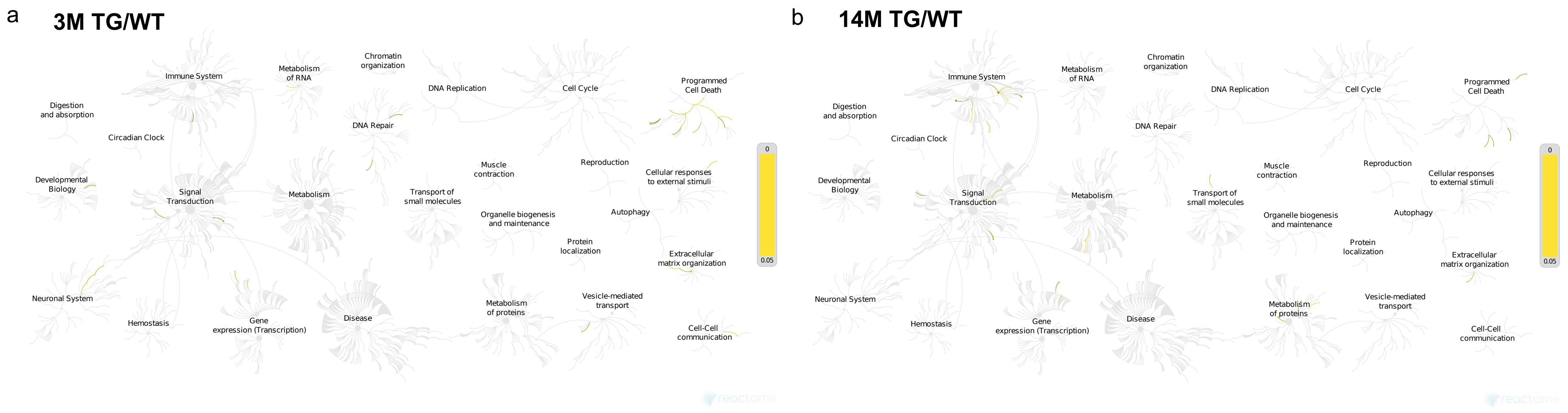
a
3M TG/WT
Immune System
Metabolism of RNA
Chromatin organization
DNA Replication
Cell Cycle
Programmed Cell Death
Digestion and absorption
Circadian Clock
DNA Repair
Developmental Biology
Signal Transduction
Metabolism
Transport of small molecules
Muscle contraction
Reproduction
Cellular responses to external stimuli
Organelle biogenesis and maintenance
Autophagy
Protein localization
Extracellular matrix organization
Neuronal System
Hemostasis
Gene expression (Transcription)
Disease
Metabolism of proteins
Vesicle-mediated transport
Cell-Cell communication
0
0.05
b
14M TG/WT
Immune System
Metabolism of RNA
Chromatin organization
DNA Replication
Cell Cycle
Programmed Cell Death
Digestion and absorption
Circadian Clock
DNA Repair
Developmental Biology
Signal Transduction
Metabolism
Transport of small molecules
Muscle contraction
Reproduction
Cellular responses to external stimuli
Organelle biogenesis and maintenance
Autophagy
Protein localization
Extracellular matrix organization
Neuronal System
Hemostasis
Gene expression (Transcription)
Disease
Metabolism of proteins
Vesicle-mediated transport
Cell-Cell communication
0
0.05

Suppl. Fig. 13

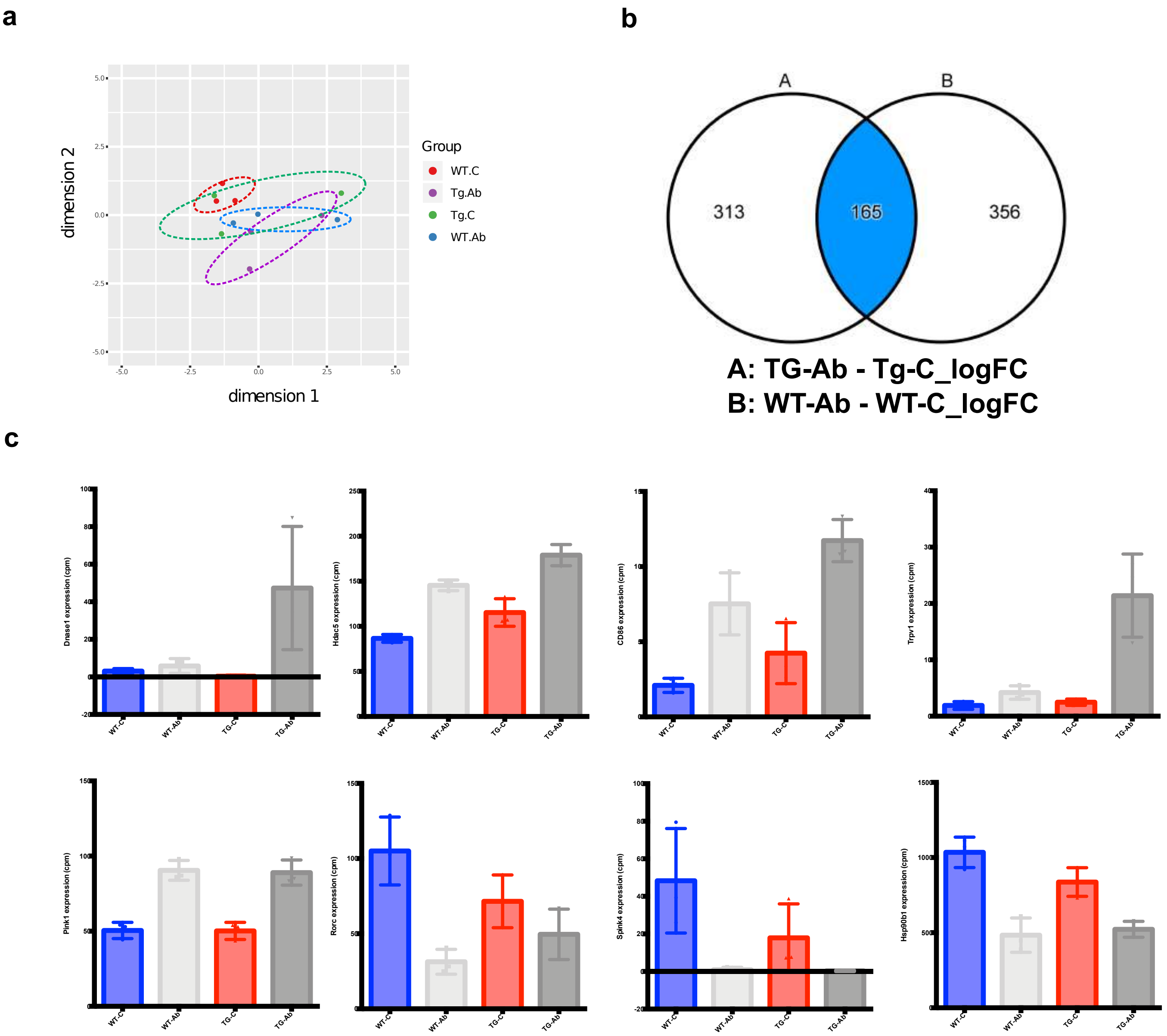
a
b
c
dimension 2
dimension 1
Group
WT.C
Tg.Ab
Tg.C
WT.Ab
A
B
313
165
356
A: TG-Ab - Tg-C_logFC
B: WT-Ab - WT-C_logFC
Dnase1 expression (cpm)
Hdac5 expression (cpm)
CD86 expression (cpm)
Trpv1 expression (cpm)
Pink1 expression (cpm)
Rorc expression (cpm)
Spink4 expression (cpm)
Hsp90b1 expression (cpm)
WT-C
WT-Ab
TG-C
TG-Ab

Suppl. Fig. 14

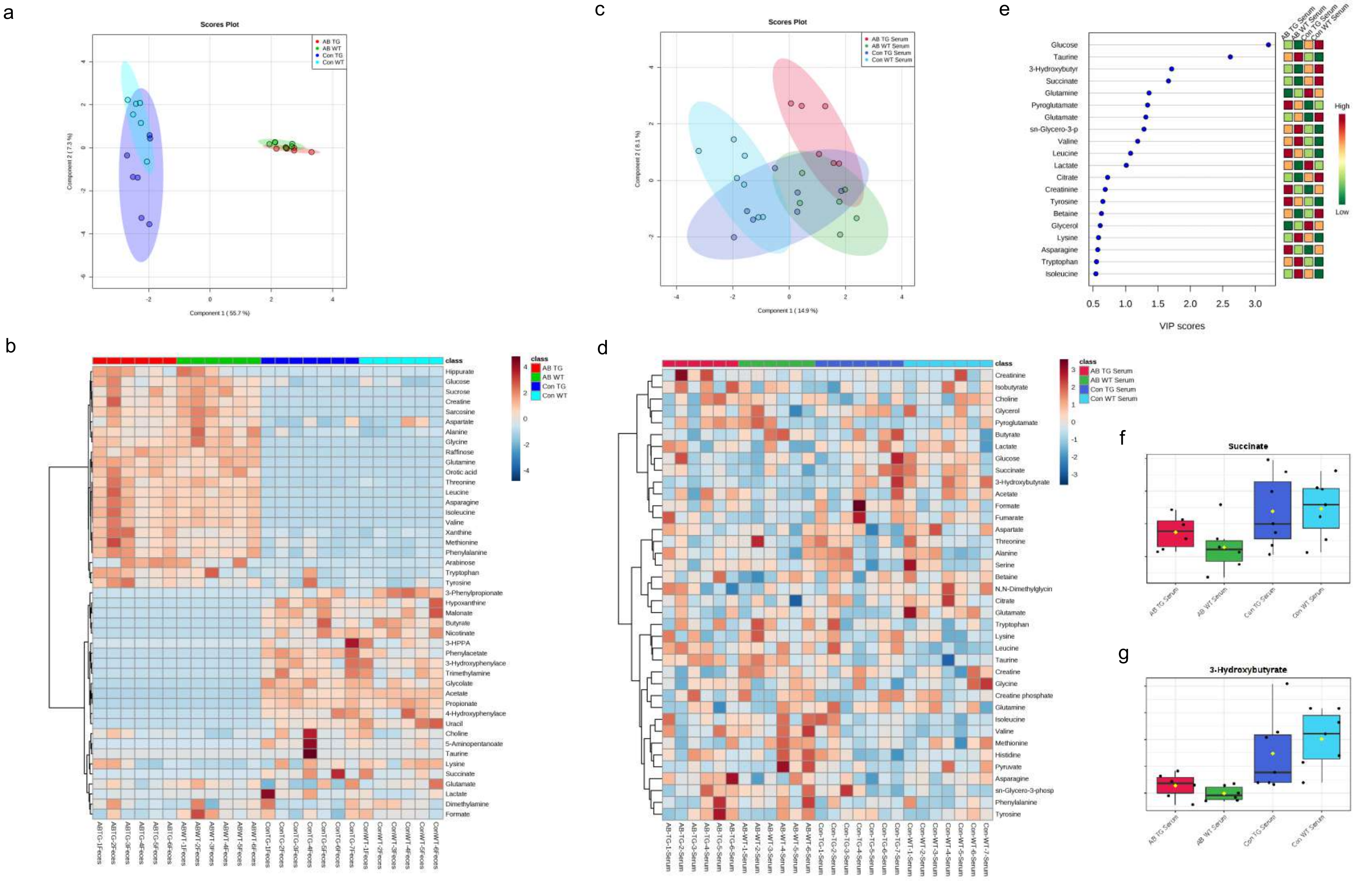
a
Scores Plot
AB TG
AB WT
Con TG
Con WT
Component 1 ( 55.7 %)
Component 2 ( 7.3 %)
b
class
AB TG
AB WT
Con TG
Con WT
Hippurate
Glucose
Sucrose
Creatine
Sarcosine
Aspartate
Alanine
Glycine
Raffinose
Glutamine
Orotic acid
Threonine
Leucine
Asparagine
Isoleucine
Valine
Xanthine
Methionine
Phenylalanine
Arabinose
Tryptophan
Tyrosine
3-Phenylpropionate
Hypoxanthine
Malonate
Butyrate
Nicotinate
3-HPPA
Phenylacetate
3-Hydroxyphenylace
Trimethylamine
Glycolate
Acetate
Propionate
4-Hydroxyphenylace
Uracil
Choline
5-Aminopentanoate
Taurine
Lysine
Succinate
Glutamate
Lactate
Dimethylamine
Formate
c
Scores Plot
AB TG Serum
AB WT Serum
Con TG Serum
Con WT Serum
Component 1 ( 14.9 %)
Component 2 ( 8.1 %)
d
class
AB TG Serum
AB WT Serum
Con TG Serum
Con WT Serum
Creatinine
Isobutyrate
Choline
Glycerol
Pyroglutamate
Butyrate
Lactate
Glucose
Succinate
3-Hydroxybutyrate
Acetate
Formate
Fumarate
Aspartate
Threonine
Alanine
Serine
Betaine
N,N-Dimethylglycin
Citrate
Glutamate
Tryptophan
Lysine
Leucine
Taurine
Creatine
Glycine
Creatine phosphate
Glutamine
Isoleucine
Valine
Methionine
Histidine
Pyruvate
Asparagine
sn-Glycero-3-phosp
Phenylalanine
Tyrosine
e
AB TG Serum
AB WT Serum
Con TG Serum
Con WT Serum
Glucose
Taurine
3-Hydroxybutyr
Succinate
Glutamine
Pyroglutamate
Glutamate
sn-Glycero-3-p
Valine
Leucine
Lactate
Citrate
Creatinine
Tyrosine
Betaine
Glycerol
Lysine
Asparagine
Tryptophan
Isoleucine
High
Low
VIP scores
f
Succinate
AB TG Serum
AB WT Serum
Con TG Serum
Con WT Serum
g
3-Hydroxybutyrate
AB TG Serum
AB WT Serum
Con TG Serum
Con WT Serum

Suppl. Fig. 15
